# ERK plays a conserved dominant role in pancreas cancer cell EMT heterogeneity driven by diverse growth factors and chemotherapies

**DOI:** 10.1101/2025.02.08.637251

**Authors:** Michelle C. Barbeau, Brooke A. Brown, Sara J. Adair, Todd W. Bauer, Matthew J. Lazzara

**Author notes:** Corresponding Author: Matthew J. Lazzara, 385 McCormick Road, Charlottesville, VA 22903. Summary: A common basis for EMT to occur in cancer cell sub-populations in response to mitogenic, differentiation-inducing, and DNA-damaging conditions has been elusive, but iterative immunofluorescence imaging and information theory modeling of cell populations revealed that ERK plays a surprisingly conserved role across diverse EMT drivers and that ERK inhibition antagonizes EMT to promote chemoresponse.

## Abstract

Epithelial-mesenchymal transition (EMT) occurs heterogeneously among malignant carcinoma cells to promote chemoresistance. Identifying the signaling pathways involved will nominate drug combinations to promote chemoresponse, but cell population-level studies are inherently fraught, and single-cell transcriptomics are limited to indirect ontology-based inferences. To understand EMT heterogeneity at a signaling protein level, we combined iterative indirect immunofluorescence imaging of pancreas cancer cells and tumors and mutual information (MI) modeling. Focusing first on MAP kinase pathways, MI predicted that cell-to-cell variation in ERK activity surprisingly dominated control of EMT heterogeneity in response to diverse growth factors and chemotherapeutics, but that JNK compensated when MEK was inhibited. Population-level models could not capture these experimentally validated MI predictions. The dominant role of ERK was predicted by MI even when analyzing seven potential EMT-regulating signaling nodes. More generally, this work provides an approach for studying highly multivariate signaling/phenotype relationships based on protein measurements in any setting.

## INTRODUCTION

The epithelial-mesenchymal transition (EMT) is a developmental program that is aberrantly reactivated in numerous carcinomas, resulting in cells that are more invasive and drug-resistant (*1*). Cells undergoing EMT lose apical-basal polarity, downregulate tight junction and adherens junction proteins (e.g., E-cadherin), and increase expression of the intermediate filament protein vimentin. These and other changes are coordinated by a well characterized set of transcription factors. The functional relevance of EMT for invasion and drug resistance has been clearly demonstrated in carcinomas including pancreatic ductal adenocarcinoma (PDAC) (*2–4*), an almost uniformly lethal cancer that is increasingly common. PDAC tumors have high EMT heterogeneity, with as many as 33 to 67 percent of ductal cells displaying mesenchymal characteristics (*2, 5*). Heterogeneity may be genetic in origin or may arise from local tumor gradients in cytokines, oxygen, or non-transformed stromal cells that signal to neoplastic cells (*6, 7*). Even in vitro, cells experiencing nominally identical conditions exhibit EMT heterogeneity (*8, 9*).

The tumor microenvironment provides a variety of routes for EMT in malignant cells. Nonmalignant cancer-associated fibroblasts and tumor-associated macrophages secrete TGFβ, ligands for receptor tyrosine kinases (e.g., EGF, FGF, HGF), interleukins (e.g., IL6, IL8, and IL10), and TNFα, which are potent activators of SMADs, MAPKs, JAK/STAT, and NFκB signaling, respectively (*10–20*). EMT is most robustly driven in carcinoma cells through cooperation of distinct signaling pathways, for example, in response to EGF and TGFβ (*21*). Other EMT-relevant signaling pathways are activated by cell-matrix interactions and tumor hypoxia (*22–25*). DNA-damaging chemotherapies, gemcitabine, and drug combinations including FOLFIRINOX also drive EMT in PDAC (*2, 26*). Collectively, these observations strongly suggest that EMT is controlled by the concerted activity of multiple diverse signaling pathways, adding an extra degree of difficulty to identifying the signaling basis for EMT heterogeneity.

The few existing computational or conceptual models of EMT heterogeneity are based primarily on the predicted effects of stochastic partitioning of EMT-suppressive factors such as GRHL2, Ovol2, miR-200 and miR-34a, which regulate EMT transcription factors including Zeb1 and Snail (*27–30*). The heavy reliance of these studies on TGFβ as the EMT agonist and lack of focus on kinase-regulated signaling pathways mapping to approved or investigational inhibitors makes their results challenging to use for the design of EMT-antagonizing therapeutic combinations that are likely to work in the complex tumor microenvironment. Kinase-regulated signaling nodes represent a good domain to search for drug targets that control PDAC EMT. Indeed, cell-to-cell variation in kinase-regulated pathway nodes drive phenotypic heterogeneity in contexts including response to chemotherapy (*31, 32*). Thus, this avenue may offer key, targetable insights into EMT regulation.

The complexity of EMT heterogeneity calls for systematic approaches for measuring the key regulatory signaling pathways and quantitatively mapping them to phenotypes. Single-cell RNA sequencing (scRNAseq) is a common high-resolution method for measuring heterogeneous phenotypes, but it relies on indirect inferences of protein activation states (*33*). Flow- and mass-cytometry require cell or tumor disaggregation, which disrupt labile phosphoproteins and involve costly or complex protocols requiring conjugated antibodies or compensation approaches (*34, 35*). To overcome these barriers, we used a multiplexed iterative imaging approach that produces single-cell protein-level measurements in fixed cells and tumors (*36*). To interpret the resulting high-dimensional datasets of signaling and phenotype measurements, we used a mutual information (MI) computational model that predicts the extent to which individual pathways or combinations inform the cell EMT state (*37*). MI models do not require *a priori* knowledge of the relationship among the variables measured. That is, unlike parametric modeling approaches such as partial least-squares regression, MI models do not assume linearity or any other algebraic form in the relationship between signal activities and phenotype. These model characteristics increase the odds of identifying real quantitative relationships between heterogeneously activated druggable signaling pathways and EMT. Using this workflow, we identified the conserved roles of the ERK and JNK pathways (among a larger set of measured signaling pathway nodes) in driving EMT in response to a diverse set of growth factors and chemotherapeutic drugs (*11, 38*). Cell culture and in vivo studies of PDAC model systems demonstrated that ERK exerts dominant control over EMT heterogeneity but that JNK compensated for impaired ERK activity when MEK was inhibited. These results can inform the design of combination therapy approaches for the treatment of PDAC that will minimize the chemoresistance that arises due to EMT.

## RESULTS

### A mutual information model nominates ERK over JNK as a key regulator of growth factor-driven EMT

To investigate the ability of cell-to-cell variation in specific signaling pathway activity levels to explain EMT heterogeneity in populations of cells under diverse treatment conditions, we first examined the issue for ERK and JNK. ERK plays important roles in growth factor-mediated EMT (*21, 39*), and ERK and JNK regulate hypoxia-driven EMT (*40*), but ways in which these pathways control EMT heterogeneity is uknown. We generated a dataset by making single-cell, protein-level measurements of EMT phenotype markers and signaling protein phosphorylation using iterative indirect immunofluorescence imaging (4i) because the total number of signals needed for even this relatively simple analysis exceeded what can be done in a single round of standard immunofluorescence imaging **(Figure 1A)** (*36*). Briefly, 4i involves repeated rounds of antibody staining and immunofluorescence imaging, with each round followed by an antibody elution step. This enables the acquisition of a potentially arbitrary number of signals using a conventional 4-channel fluorescence microscope and a limited number of distinct species sources for unconjugated primary antibodies, while retaining the benefit of signal amplification enabled by indirect immunofluorescence. For each antibody in the panel, we confirmed its efficient elution from the sample, tested signal preservation across rounds of staining and elution, and validated its utility for detecting pathway response to growth factors **(SuppFigure S1A-C).** To study the basis for EMT heterogeneity in terms of ERK and JNK pathway activity, we used a panel of five antibodies (plus a DNA stain) across three rounds of 4i for a total of 6 channels **(Figure 1B; SuppTable S1).** ERK1/2 phosphorylation (Thr202/Tyr204) was measured as a proxy for ERK activity, and c-Jun phosphorylation (Ser63) was measured as a proxy for JNK activity (*40–42*). Total ERK was measured to provide a signal for cell segmentation in image analysis. Antibodies for E-cadherin and vimentin were included to assess cell EMT status.

**Figure 1.**
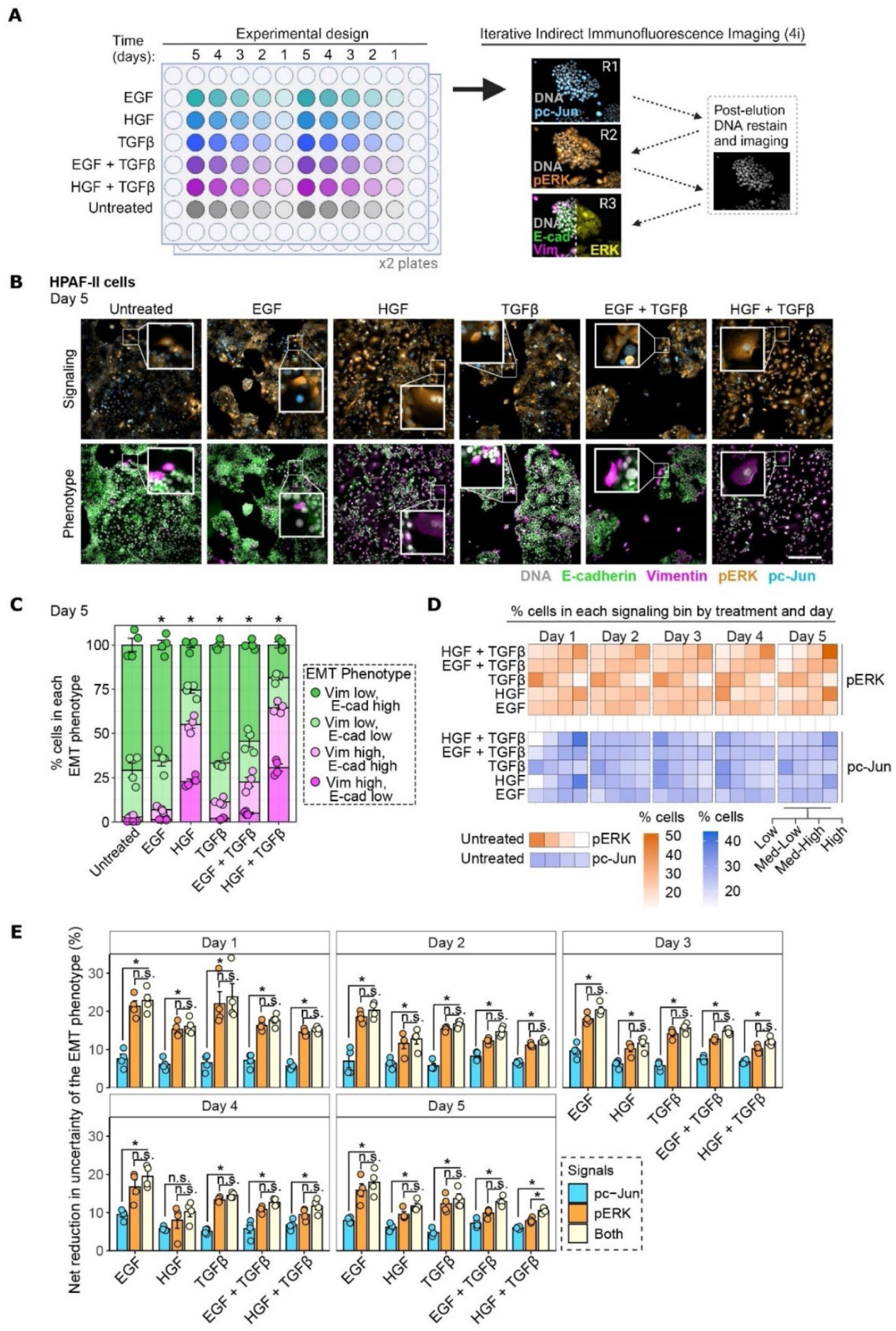
A mutual information model based on immunofluorescence measurements predicts that ERK is dominant over JNK for determining EMT heterogeneity. **A)** Schematic of the 96-well format used for 4i experiments and the iterative indirect immunofluorescence imaging (4i) protocol used after plate fixation. The details shown are for the growth factor-mediated EMT experiment conducted and analyzed in the remainder of this figure. R = Round. **B)** HPAF-II cells were treated for 1-5 days with or without 50 ng/mL EGF, 50 ng/mL HGF, 10 ng/mL TGFβ, EGF+TGFβ or HGF+TGFβ. Representative 4i images are shown for the indicated proteins at the day-5 timepoint, n = 4. **C)** For the experiment described in (B) at the day-5 timepoint, mosaic plots are shown displaying percentages of cells in each EMT phenotype bin. n = 4, Chi-squared test for comparisons against untreated control. Vim = vimentin and E-cad = E-cadherin. **D)** For the experiment described in (B), the heatmaps show percentages of cells in each pERK (top) or pc-Jun (bottom) signaling bin. **E)** For the experiment described in (B), the net percent reduction in EMT phenotype uncertainty based on knowing pERK, pc-Jun, or both signals is plotted. n = 4, one-way ANOVA with Tukey’s multiple comparisons test. For all panels, *p < 0.05, and error bars represent standard error of the mean. Scale bar indicates 300 μm.

For the first dataset, we used HPAF-II human pancreas cancer cells, which exhibit baseline epithelial characteristics and an additive or synergistic EMT response to diverse growth factors(*21, 40*). EMT was induced with growth factors for three different receptor kinases (EGF:EGFR, HGF:MET, and TGFβ:TGFβR) known to drive EMT and with the combinations EGF+TGFβ and HGF+TGFβ (*43*). Treatment time ranged from 1-5 days to increase opportunities to observe co-occurring protein phosphorylation events and phenotypic marker changes. EMT occurred heterogeneously in every condition, which was visually apparent by the variable retention of E-cadherin and incomplete expression of vimentin among cells. With the development of an MI model in mind, the degree of EMT heterogeneity was quantified by binning cells into phenotype states based on high or low expression of membranous E-cadherin or whole-cell vimentin expression. Cutoffs were defined based on deviation from the untreated baseline epithelial condition (*Materials and Methods*). This created a single measure for EMT state informed by the combination of high/low states of two independent protein measurements, resulting in four distinct “EMT phenotype” bins across experimental conditions **(Figure 1C, SuppFigure S2).** Stacked mosaic plots of phenotype such as the one in Figure 1C are shown throughout with the most epithelial state at top and most mesenchymal state at bottom. The two intermediate phenotypes are positioned to create contiguous low- and high-vimentin groupings of states. This facilitates interpretation of some results based on changes in the total fraction of vimentin-high cells, which we favor as a clinically relevant marker for PDAC patient prognosis (*44*).

To relate signaling activation to EMT changes, protein phosphorylation signals were discretized into bins referred to as “signaling states.” The optimal number of signaling states (across growth factors and days) was determined using Gaussian mixture models of pERK and pc-Jun and computing the Bayesian information criterion **(SuppFigure S3A,B**; *Materials and Methods*). There was no benefit of exceeding four discrete signaling states. Thus, the bounds for the states of pERK and pc-Jun were chosen to create four evenly percentiled bins capturing low, medium-low, medium-high, or high levels of phosphorylation of each protein, considering all times and conditions. Note that the four phenotype states were chosen on a different basis and that the difference in binning approaches for EMT phenotype and signaling states creates a situation where the MI models assessed how graded changes in signaling promote more switch-like changes in EMT. Once set for building a model, the cutoffs for all bins were invariant. As shown in **Figure 1D**, the distribution of cells in each signaling state varied by treatment condition and time. For example, there was clearly increased occupancy of the high bin for pERK and pc-Jun for HGF-containing treatments and increased occupancy of the low bin for treatment with TGFβ alone.

After converting the continuous metrics extracted from cell images to the discrete phenotype and signaling states metrics, the joint and marginal probabilities for each binned state and combinations of signaling and EMT phenotype state were calculated and used to build an MI model quantifying the extent to which the signaling states explained intrapopulation heterogeneity in EMT phenotype for each condition (*Materials and Methods*). In the MI model generated, pERK consistently provided more information about the EMT phenotype than did pc-Jun **(Figure 1E)**. In most cases, pc-Jun provided only redundant information – i.e., the MI of pc-Jun and pERK together was not significantly greater than that for pERK alone. Put another way, the association between signaling activation and EMT phenotype did not increase even if c-Jun was also phosphorylated. Thus, variation in ERK activity was predicted to be a more important determinant of EMT heterogeneity than variation in JNK activity, though many cells demonstrated a clear increase in pc-Jun. Results shown throughout subsequent figures demonstrate that this model inference was generalizable to other cell backgrounds and EMT agonists.

To extend the analysis to additional backgrounds, we tested baseline-epithelial murine 7160c2 cells derived from a KPCY (Kras^LSL-G12D^, p53^loxP/+ or^ ^LSL-R172H^, Pdx1-Cre, Rosa26^LSL-YFP^) mouse model of PDAC. Two-day treatment with EGF+TGFβ or HGF+TGFβ induced robust phosphosignals and substantial EMT **(Figure 2A-C)**, with significantly more EMT MI for pERK than pc-Jun **(Figure 2D)**, consistent with the model for HPAF-II cells. We also analyzed PDX 395 cells, which were isolated from a human PDAC patient-derived xenograft (PDX) and exhibited baseline EMT heterogeneity under standard culture conditions **(Figure 2E-F)**. A model based on images of PDX 395 cells without addition of any EMT agonists demonstrated significantly more EMT MI for pERK than pc-Jun **(Figure 2G)**, consistent with inferences from growth factor-induced EMT in HPAF-II and 7160c2 cell lines. In aggregate, these results suggest a generalizable conclusion that ERK is a more important determinant of EMT heterogeneity than JNK is in determining EMT at baseline and in response to growth factors.

**Figure 2.**
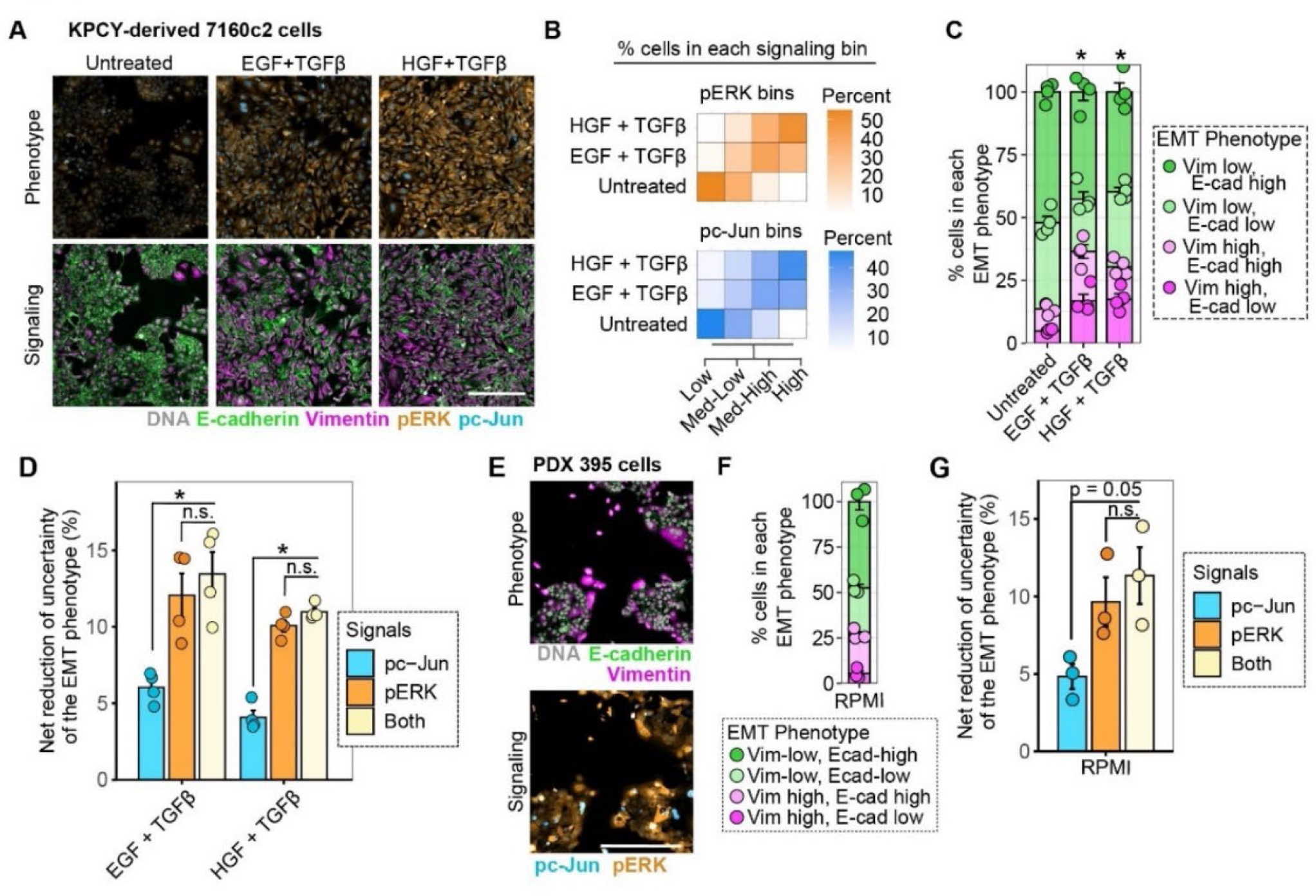
Mutual information models point to ERK as controlling EMT heterogeneity in multiple cell settings. **A)** KPCY-derived 7160c2 cells were treated for 2 days with or without 50 ng/mL EGF + 10 ng/mL TGFβ or 50 ng/mL HGF + 10 ng/mL TGFβ. Representative 4i images are shown for the indicated proteins, n = 4. **B)** For the experiment described in (A), heatmaps are shown of the percent of cells in each pc-Jun (top) or pERK (bottom) signaling bin. **C)** For the experiment described in (A), the mosaic plot displays percentages of cells in each EMT phenotype bin. n = 4, Chi-squared test for comparisons against untreated control. **D)** For the experiment described in (A), the net percent reduction in uncertainty of the EMT phenotype based on knowing pERK, pc-Jun, or both signals is plotted. n = 4, one-way ANOVA with Tukey’s multiple comparisons test. **E)** PDX 395 cells were cultured under normal conditions. Representative 4i images are shown for the indicated proteins, n = 3. **F)** For the experiment described in (E), the mosaic plot displays the percentage of cells in each EMT phenotype bin, n = 3. **G)** For the experiment described in (E), the net percent reduction in EMT phenotype, based on knowing pERK, pc-Jun, or both signals, is plotted. n = 3, one-way ANOVA with Tukey’s multiple comparisons test. For all panels *p < 0.05, and error bars represent standard error of the mean. Scale bar indicates 300 μm.

### Targeted pathway antagonism confirms the predicted role of ERK and reveals a compensatory role for JNK in growth factor-driven EMT

To test the predicted preferential importance of ERK over JNK for EMT in HPAF-II cells, we assessed the relative effects of inhibitors of MEK (trametinib) and/or JNK (SP600125) on EMT induced by EGF+TGFβ or HGF+TGFβ **(Figure 3A)**. SP600125 was used at a concentration that suppressed pc-Jun without suppressing pERK **(SuppFigure S4A-C)**. Trametinib was used at a concentration that consistently suppressed pERK but also led to an unexpected increase in pc-Jun at later timepoints **(SuppFigure S4D-F)**. We examined cells after four days because the unexpected pc-Jun signal was evident by that time. EMT was reduced by both inhibitors, but MEK inhibition had a significantly greater effect than JNK inhibition **(Figure 3B)**, consistent with the MI model prediction. To examine the role of the late pc-Jun signal in MEK-inhibited cells, MI was computed between pc-Jun and EMT in the presence of trametinib on day 4. In MEK-inhibited cells, the elevated pc-Jun signal corresponded with a significant increase in pc-Jun MI for EMT, suggesting that JNK activation compensated for MEK/ERK signaling to promote EMT when MEK/ERK was inhibited **(Figure 3C-D).**

**Figure 3.**
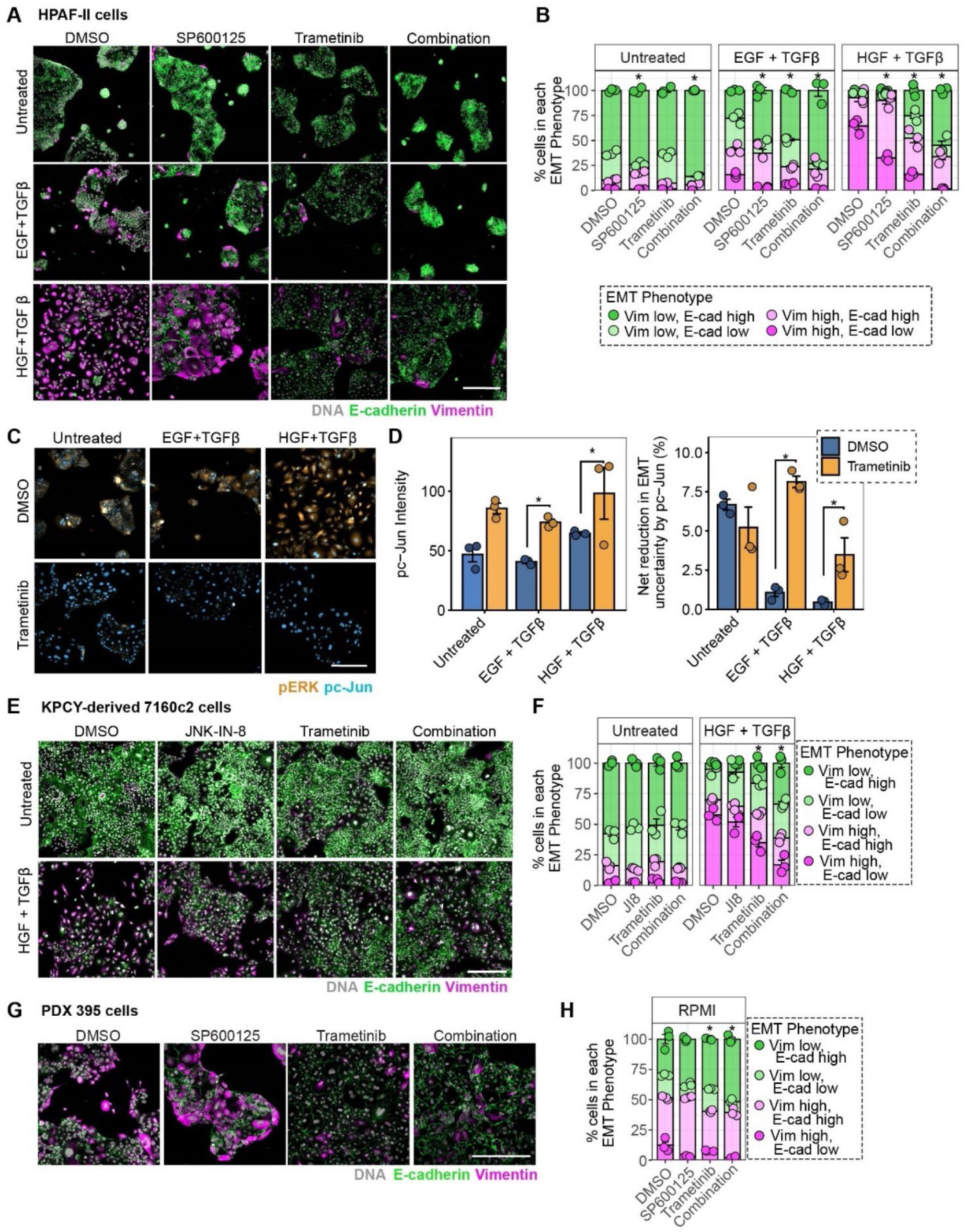
Inhibition and knockdown studies confirm a dominant role of ERK in EMT heterogeneity and a compensatory role for JNK when MEK is inhibited. **A)** HPAF-II cells were treated with or without 50 ng/mL EGF + 10 ng/mL TGFβ or 50 ng/mL HGF + 10 ng/mL TGFβ and DMSO, 10 μM SP600125 (JNKi), 200 nM trametinib (MEKi), or trametinib + SP600125. Representative 4i images are shown for the indicated proteins, n = 3. **B)** For the experiment described in (A), the mosaic plot displays percentages of cells in each EMT phenotype bin. n = 3, Chi-squared test. **C)** For the experiment described in (A), representative 4i images of indicated proteins are shown, n = 3. **D)** For the experiment described in (A), intensity of pc-Jun in the presence or absence of trametinib (left) and mutual information model of pc-Jun with EMT phenotype (right) +/- trametinib are plotted, n = 3, one-way ANOVA with Tukey’s multiple comparisons test. **E)** KPCY-derived 7160c2 cells were treated with or without 50 ng/mL HGF + 10 ng/mL TGFβ and DMSO, 5 μM JNK-IN-8 (JI8), 2 nM trametinib, or both inhibitors for 3 days. Representative 4i images of the indicated proteins are shown, n = 3. **F)** For the experiment described in (E), the mosaic plot displays the percentage of cells in each EMT phenotype bin. n = 3, Chi-squared test. **G)** PDX 395 cells were treated with DMSO, 10 μM SP600125, 20 nM trametinib, or trametinib + SP600125 for 3 days, n = 3. **H)** For the experiment described in (G), the mosaic plot displays the percentages of cells in each EMT phenotype bin. n = 3, Chi-square test. For all panels *p < 0.05, and error bars represent standard error of the mean. Scale bars indicate 300 μm.

To validate the model in another cell background, 7160c2 cells were treated with HGF+TGFβ with or without trametinib and the JNK inhibitor JNK-IN-8 **(Figure 3E),** using concentrations that inhibited their targets **(SuppFigure S4G-H)**. JNK-IN-8 was chosen for its superior ability to inhibit pc-Jun in 7160c2 cells compared with SP600125 (**SuppFigure S4I)**. Again, MEK inhibition more effectively antagonized EMT than did JNK inhibition **(Figure 3F).** PDX 395 cells were also treated with trametinib and/or SP600125 at concentrations shown to inhibit their targets **(SuppFigure S4J,K)**. Once again, EMT was antagonized significantly more by MEK inhibition than by JNK inhibition **(Figure 3G,H)**.

To test the MI model using an alternative to inhibition, siRNA-mediated knockdown was used in HPAF-II cells treated with growth factors. siRNA-mediated knockdown of ERK1/2 and c-Jun was efficient, although a perplexing apparent pc-Jun increase occurred in cells not treated with growth factors that did not affect our interpretation of results **(SuppFigure S5A-F)**. In growth factor-treated HPAF-II cells, ERK knockdown reduced the fraction of vimentin-positive cells and prevented loss of E-cadherin to a much greater degree than did c-Jun knockdown **(Figure S5G-H)**, consistent with MI model predictions. Furthermore, the natural heterogeneity in ERK knockdown among cells, which can be observed in the total ERK immunofluorescence signal **(Figure S5I)**, provided another opportunity to test the MI model. Specifically, we hypothesized that the relationship between phosphosignals and phenotype would be preserved even when the fraction of cells exhibiting high-pERK levels was altered by ERK knockdown. Indeed, MI calculations suggested a consistently greater role for pERK in regulating EMT heterogeneity across all siRNA conditions **(Figure S5J)**. The aggregated results using inhibitors and siRNAs support our hypothesis that ERK dominates the regulation of EMT under the cytokine-treated or baseline conditions but that JNK compensates for ERK to drive EMT in the presence of a strong MEK inhibitor.

### ERK is also preferentially important over JNK for chemotherapy-induced EMT

In search of a condition where JNK (a stress-activated kinase) played the dominant role over ERK, we turned to chemotherapy as a potential EMT driver (*45*). We chose 5-fluorouracil and gemcitabine, which are both used to treat pancreas cancer, and selected concentrations and timing resulting in minimal cell death. In HPAF-II cells, 5-fluorouracil or gemcitabine significantly increased the number of vimentin-positive cells, and gemcitabine significantly reduced membranous E-cadherin **(Figure 4A,B)**. Vimentin expression was comparable to 48-hr EGF+TGFβ treatment in HPAF-II cells, and gemcitabine-mediated E-cadherin loss was stronger than that observed for any growth factor condition. These protein-level changes were accompanied by robust increases in mRNAs for the EMT transcription factors *SLUG, SNAIL*, and *ZEB 1* **(SuppFigure S6 A,B)**, confirming that a *bona fide* EMT occurred. Even though the induction of pc-Jun was more visually impressive than that of pERK in the immunofluorescence imaging, MI analysis surprisingly suggested a greater importance of pERK in determining the heterogeneous EMT phenotype **(Figure 4C)**. We repeated the experiment using PDX 395 cells, which again showed significant EMT in response to chemotherapy **(Figure 4D,E)** that had higher MI with pERK than pc-Jun **(SuppFigure S6C)**. The experiment was repeated with 7160c2 cells and gemcitabine. A 24-hr gemcitabine pulse (rather than continuous treatment) was used for 7160c2 cells because they displayed high sensitivity to the drug, presumably owing to their very high proliferation rate. 7160c2 cells displayed significant induction of EMT in response to gemcitabine **(Figure 4F)**, which was again better explained by pERK than pc-Jun according to the MI model for that cell line **(SuppFigure S6D,E)**. HPAF-II and PDX 395 cells were used to test the MI models. Consistent with model predictions, in HPAF-II cells treated with gemcitabine or 5-fluorouracil, EMT was more substantially inhibited by co-treatment with trametinib than with SP600125 **(Figure 4G-H)**. In PDX 395 cells treated with 5-fluorouracil, vimentin expression was better inhibited by cotreatment with trametinib than with SP600125 **(SuppFig S6 F,G)**. Taken together, the data showed that ERK also plays a dominant role over JNK in regulating EMT in response to chemotherapeutics.

**Figure 4.**
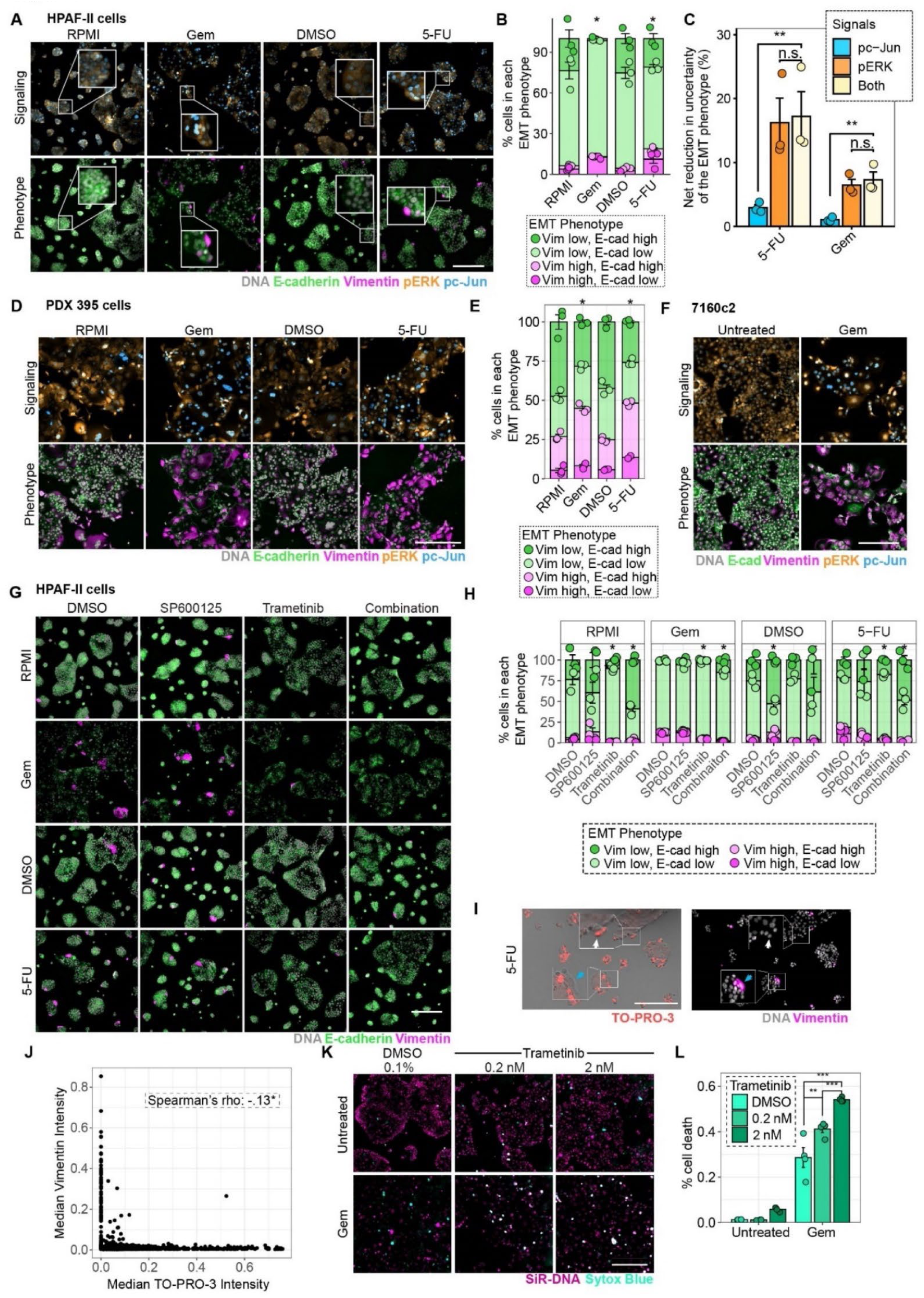
ERK plays a dominant role over JNK in determining EMT heterogeneity in response to chemotherapy. **A)** HPAF-II cells were treated with or without 10 ng/mL gemcitabine (Gem) or 10 ng/mL 5-fluorouracil (5-FU) for 2 days. RPMI and DMSO were used as controls for gemcitabine or 5-FU, respectively. Representative 4i images for the indicated proteins are shown, n = 3. **B)** For the experiment described in (A), the mosaic plot displays percentages of cells in each EMT phenotype bin. n = 3, Chi-squared test. **C)** For the experiment described in (A), the net percent reduction in uncertainty of the EMT phenotype, based on knowing pERK, pc-Jun, or both signals, is plotted. n = 3, one-way ANOVA with Tukey’s multiple comparisons test. **D)** PDX 395 were cells treated with or without 10 ng/mL gemcitabine or 10 ng/mL 5-FU for 3 days. RPMI and DMSO were used as controls for gemcitabine or 5-FU, respectively. Representative 4i images for the indicated proteins, n = 3. **E)** For the experiment described in (D), the mosaic plot displays percentages of cells in each EMT phenotype bin. n = 3, Chi-squared test. **F)** KPCY-derived 7160c2 were pulsed with or without 10 ng/mL gemcitabine for 24 hr and measured for signaling and EMT markers at 48 hr. Representative 4i images of the indicated proteins, n = 3. **G)** HPAF-II cells were treated with or without 10 ng/mL 5-FU or 10 ng/mL gemcitabine and additionally treated with or without 10 μM SP600125, 200 nM trametinib, or both for 2 days. RPMI and DMSO were used as controls for gemcitabine or 5-FU, respectively. Representative 4i images of the indicated proteins, n = 3. **H)** For the experiment described in (G), the mosaic plot displays percentages of cells in each EMT phenotype bin. n = 3, Chi-squared test. **I)** HPAF-II cells were treated with 10 ng/mL 5-FU for 48 hr followed by a washout with complete media for 72 hr. Representative live-cell fluorescence image of live-cell-impermeant TO-PRO-3 (left) and paired fixed immunofluorescence image of vimentin (right), n = 3. White arrow indicates a representative TO-PRO-3 positive, vimentin-negative cell. Blue arrow indicates a representative TO-PRO-3-negative, vimentin-positive cell. **J)** For the experiment described in (I), TO-PRO-3 intensity versus vimentin intensity is plotted. n = 610 cells pooled across 3 replicates, Spearman’s Rank Correlation. **K)** HPAF-II cells were treated with or without 1 ng/mL gemcitabine and with or without trametinib (0.2 nM or 2 nM) for 3 days. Representative live-cell images of the live-cell-permeant SiR DNA stain and live-cell impermeant Sytox Blue stain, n = 4. **L)** For the experiment described in (K), percent Sytox Blue-positive cells is plotted. n = 4, one-way ANOVA with Tukey’s multiple comparisons test. For all panels *p < 0.05, and error bars represent standard error of the mean. Scale bars indicate 300 μm.

To test if chemotherapy-induced EMT conferred a cell survival advantage, HPAF-II cells were pulsed with chemotherapeutics for 48 hr followed by 72 hr of recovery. Cells were then stained with TO-PRO-3 (permeability indicates death), imaged, and then fixed and stained for vimentin **(SuppFigure S6H)**. We observed a significant negative correlation between cell death and vimentin expression in the 5-FU-treated condition **(Figure 4I,J)**. Importantly, analyzing only TO-PRO-3 negative cells, there was a significant number of mesenchymal cells that survived 48 hr of gemcitabine or 5-FU-treatment followed by an additional 72 hr **(SuppFigure S6I)**, demonstrating that chemotherapy-induced EMT persisted past the timepoints used in Figure 4I,J. That is, the mesenchymal conversion is not a cell state that occurs along the way to inevitable cell death.

To test the hypothesis that antagonizing chemotherapy-induced EMT will increase chemosensitivity, HPAF-II cells were treated with gemcitabine and concentrations of trametinib that inhibited pERK and gemcitabine-induced mesenchymal traits **(SuppFigure S6J-L)**. Cells were stained with SiR-DNA to track live-cell nuclei and Sytox Blue to track death. Cell death was significantly augmented by co-treatment with trametinib and gemcitabine compared to gemcitabine alone **(Figure 4K,L)**. Thus, MEK inhibition may be an effective strategy for targeting cells that undergo a mesenchymal transition to survive chemotherapy.

### A cell population-level modeling misidentifies JNK as the dominant EMT-regulating pathway

To understand if the relative importance of pERK and pc-Jun for driving EMT could have been correctly inferred using population-level measurements, we developed an alternative model based on western blotting. HPAF-II cells were treated with EGF, HGF, TGFβ, EGF+TGFβ, or HGF+TGFβ. Cells were lysed at 8-hr and 5-day timepoints to measure signaling pathway nodes and EMT phenotype markers, respectively. The separation of time points was necessary because 5 days provided sufficient time for EMT to occur, but 8 hr was the latest timepoint where we could detect robust differences in signaling among conditions by blotting. The unavoidable differences in times for capturing signals and phenotypes added another difference to the population-level analysis performed versus the single-cell analyses performed for MI models.

Blot images demonstrated that EMT markers (E-cadherin and vimentin) and phosphosignals (pERK and pc-Jun) were induced to different extents among the treatment conditions **(Figure 5A-F)**. EMT was scored based on standard-scaled measurements of E-cadherin subtracted from standard-scaled measurements of vimentin. We removed one data point deemed to have undue influence based on an analysis of residuals versus leverage and checked assumptions for linear regression by inspecting plots of residuals versus fitted values, normal Q-Q, and scale-location (*Materials and Methods*) **(SuppFigure S7A-D)** (*46*). A multiple linear regression model of standard-scaled measurements with EMT score as the dependent variable and pERK and pc-Jun as the independent variables yielded an overall R^2^ = 0.87 **(Figure 5G)**. Partial regressions of each variable with EMT, which effectively hold the other variable constant, suggested that pc-Jun was more significantly predictive of the EMT phenotype than pERK **(Figure 5H-I)**, a result that stands in stark contrast to inferences from the MI models. Inferences from this population-level analysis would suggest that pc-Jun and pERK cooperated to drive EMT but that pc-Jun was more important for determining the phenotype, a conclusion that is obviously inconsistent with inferences from the experimentally validated MI model. Furthermore, a population-level model of pooled immunofluorescence data was unable to reveal significant associations with EMT from either pERK or pc-Jun **(SuppFigure S7E-F).** Overall, this analysis demonstrates the dangers of relying on population-level measurements to decipher the signaling basis for heterogeneous phenotypes and highlights the need for single-cell analyses such as those developed in this study.

**Figure 5.**
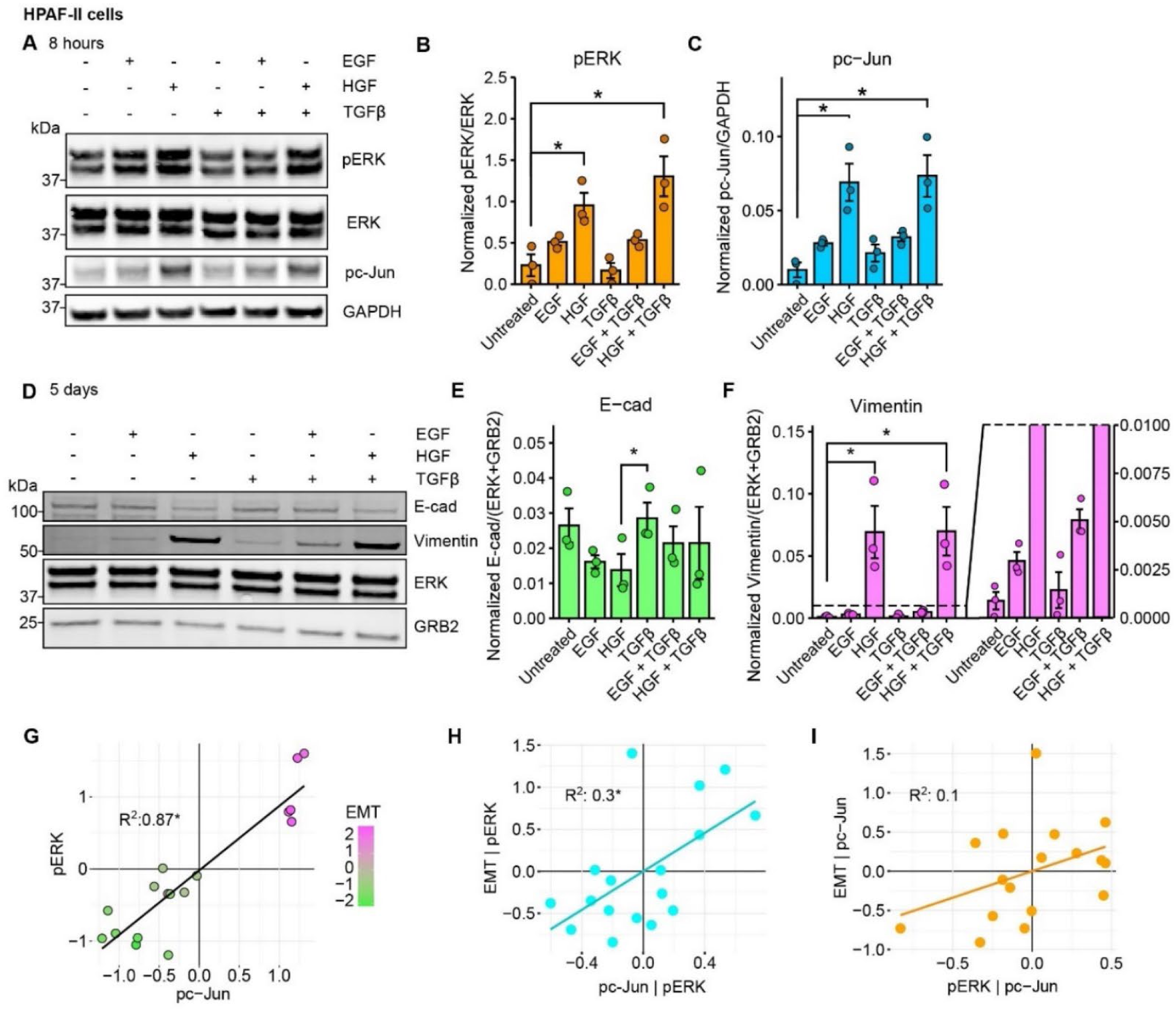
A population-level model incorrectly nominates JNK over ERK as the dominant regulator of EMT heterogeneity. HPAF-II cells were treated with or without 50 ng/mL HGF, 50 ng/mL EGF, 10 ng/mL TGFβ, EGF+TGFβ, or HGF+TGFβ and lysed 8 hr after treatment for signaling pathway analysis **(A-C)** or 5 days after treatment for EMT phenotype analysis **(D-F)**. **A)** Immunoblotting was performed for the indicated proteins. Blot image representative of n = 3. **B)** pERK intensity normalized by total ERK intensity was then median-normalized for each replicate and plotted. n = 3, one-way ANOVA with Tukey’s multiple comparisons test**. C)** pc-Jun intensity normalized by GAPDH intensity was median-normalized for each replicate and plotted. n = 3, one-way ANOVA with Tukey’s multiple comparisons test. **D)** Immunoblotting was performed for the indicated proteins. Blot image representative of n = 3. **E)** E-cadherin (E-cad) intensity normalized by total ERK and GRB2 intensities was then median-normalized and plotted. n = 3, one-way ANOVA with Tukey’s multiple comparisons test. **F)** Vimentin intensity normalized by total ERK and GRB2 intensities was then median-normalized and plotted. n = 3, one-way ANOVA with Tukey’s multiple comparisons test. **G)** GAPDH-normalized, standard-scaled pc-Jun versus pERK immunoblot measurements are plotted. Colors indicate relative EMT score as described in *Materials and Methods*. n = 3, multiple linear regression. **H)** Partial correlations of pc-Jun and EMT holding pERK constant and **I)** pERK and EMT holding pc-Jun constant. n = 3, linear least squares regression. For all panels bar plots represent the mean of the replicates, *p < 0.05, and error bars represent standard error of the mean.

### A model based on expanded signaling pathway coverage supports the importance of ERK for growth factor- and chemotherapy-driven EMT

Our analysis to this point was premised on the notion that ERK and JNK are primary controllers of EMT heterogeneity. This notion is well-grounded in the literature, but other pathways that are less well studied may also cooperate in growth factor- and chemotherapy-mediated EMT. To expand our analysis of multivariate signaling control of EMT heterogeneity, we extended the 4i antibody panel to cover a total of seven signaling nodes: whole-cell measurements for pSRC Tyr419, pp38 Thr180/Tyr182, pc-Jun Ser 73, and pERK Thr202/Tyr204, and nuclear-to-cytoplasmic ratios for SMAD2/3, STAT3, and NFκB (p65 subunit). This panel covers MAPKs, canonical EMT drivers (SMADs), and two signaling nodes reported to be important for EMT in contexts not directly considered here, STAT3 and NFκB (*47, 48*). EMT was induced using EGF+TGFβ, gemcitabine, or 5-fluorouracil. Treating HPAF-II cells with each of these agonists for 48 hr and using the full 4i panel revealed a wide range of activation states and EMT phenotypes **(Figure 6A-C; SuppTable S2).** Gemcitabine and 5-fluorouracil displayed the highest activation of the 7 pathways overall, though the EGF+TGFβ treatment condition also activated these pathways above baseline.

**Figure 6.**
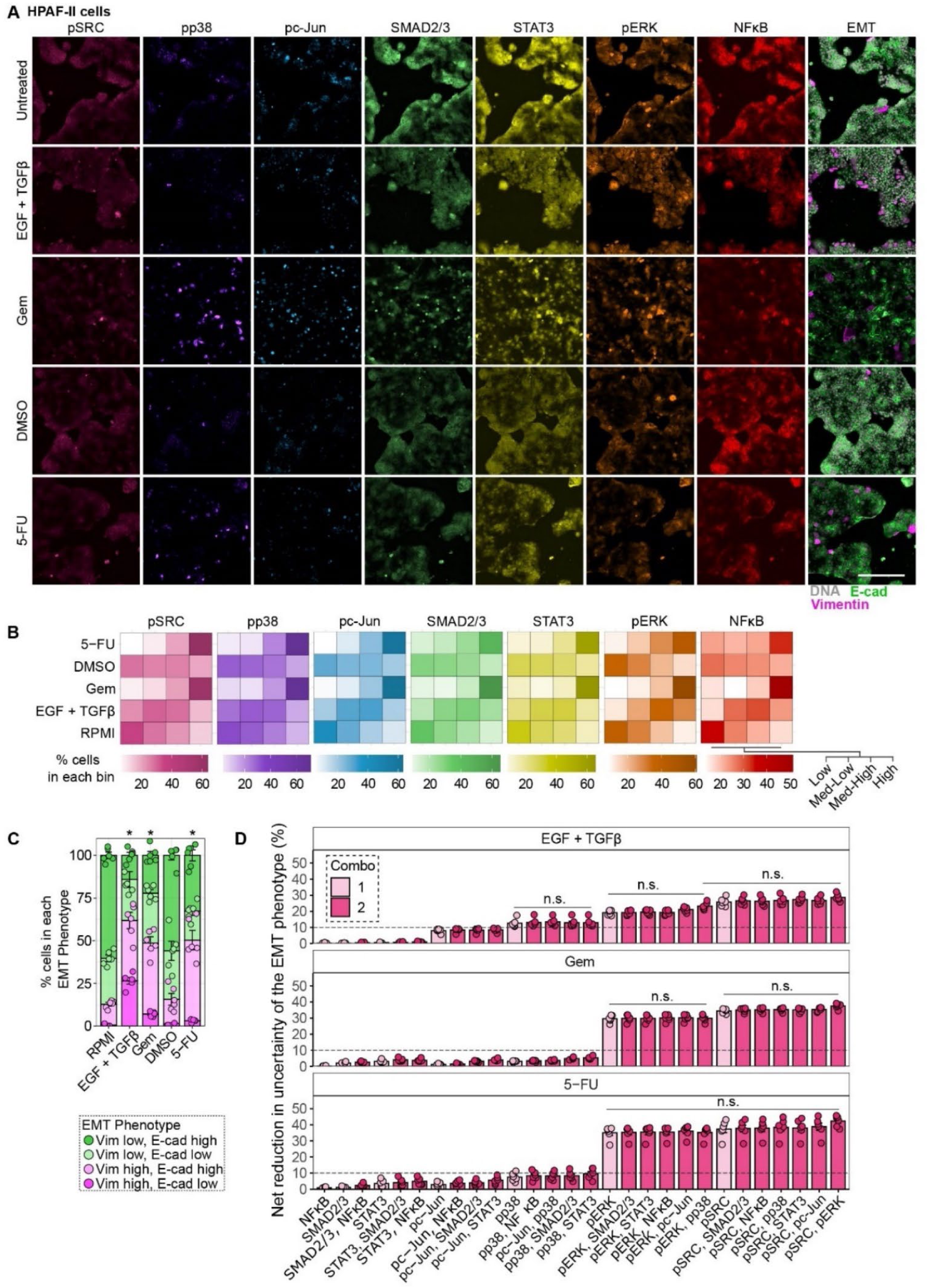
An extended multivariate mutual information model upholds the primary role of ERK in explaining EMT heterogeneity. **A)** HPAF-II cells were treated with complete media (untreated control), 10 ng/mL TGFβ + 50 ng/mL EGF,10 ng/mL gemcitabine (Gem), 10 ng/mL 5-fluorouracil (5-FU), or DMSO (control for 5-FU) for 48 hr. Representative 4i images of 7 pathway markers (pSRC, pp38, pc-Jun, SMAD2/3, STAT3, pERK, and NFκB) and 2 phenotype markers (E-cadherin and Vimentin) are shown, n = 6. **B)** For the experiment described in (A), the heatmaps show percentages of cells in each signaling pathway bin across treatment conditions, n = 6. **C)** For the experiment described in (A), the mosaic plot displays percentages of cells in each EMT phenotype bin. n = 6, Chi-squared test, *p < 0.05. **D)** For the experiment described in (A), the net percent reduction in uncertainty of the EMT phenotype from the indicated signals or combinations of signals is plotted. n = 6, one-way ANOVA with Tukey’s multiple comparisons test. Horizontal lines over vertical bars indicate that within-group comparisons are p ≥ 0.01, and comparisons with any entry outside the grouping are p < 0.01. Significance comparisons are displayed on for reductions of uncertainty ≥ 10 percent, with that threshold indicated by the dashed line. For all panels error bars represent standard error of the mean. Scale bar indicates 300 μm.

An MI model based on the data identified pERK and pSRC as playing dominant roles in determining EMT heterogeneity, with no substantial increase in MI from knowing additional signals **(Figure 6D)**. SRC family kinases (SFKs) work upstream to activate ERK and JNK and promote EMT(*40*), which may explain why pSRC and pERK each provided substantial reduction of EMT uncertainty independently without any significant benefit from knowing both. Notably, pERK and pSRC were easily the two most important individual variables in the MI model. Extending the model to three and four signaling combinations did not significantly reduce EMT uncertainty **(SuppFigure S8A-B),** which may be due to limited availability of cells populating all the higher-order signaling state combinations. Furthermore, sub-sampling increasingly unique combinations of signaling states increases the likelihood that the signaling combination only has one associated EMT phenotype, which would increase the probability of a higher MI simply by chance, as seen in the increase in MI model of the null distribution **(SuppFigure S8A-B)**. Nevertheless, the expanded MI model confirmed the importance of pERK. Interestingly, information from individual nodes including pc-Jun and pp38 was more explanatory of EMT in the EGF + TGFβ treatment than for the chemotherapy conditions.

### A mutual information model of pancreas tumor tissues reveals an in vivo shift from ERK to JNK control of EMT in response to MEK inhibition

To test the MI approach in the intact PDAC microenvironment, where a diverse set of EMT drivers is generally present, we used a 4i staining approach in PDX 395 tumors treated with or without the MEK inhibitor selumetinib **(Figure 7A; SuppTable S3).** Thresholding for cells staining positive for human COXIV to exclude murine fibroblasts from the analysis, we observed a significant decrease in vimentin-positive EMT phenotypes in the selumetinib-treated tumors **(Figure 7B)**. We then calculated MI between pERK or pc-Jun and EMT phenotype in each of the control or selumetinib-treated tumors. Due to significant inter-tumor variability, MI models were developed for individual tumors. Signaling- and phenotype-binning were performed for each tumor, and MI results were compared with a shuffled null distribution of signaling and phenotype data. We randomly sub-sampled the true and null distributions to (as described in *Material and Methods*) to create MI distributions and calculate 95% confidence intervals. Three of the four vehicle-treated control tumors showed significantly higher EMT MI for pERK than for pc-Jun, as indicated by non-overlapping confidence intervals **(Figure 7C)**. Conversely, two of the three selumetinib-treated tumors showed higher or equal reduction of uncertainty from pc-Jun than from pERK **(Figure 7D)**. The MEK-inhibited tumor that showed significantly higher pERK reduction of uncertainty also showed the highest amount of normalized pERK, suggesting that a failure to inhibit MEK substantially may explain pERK importance for explaining EMT **(Figure 7E)**. These data support a model wherein the drivers of EMT in the untreated tumors relied most heavily on ERK for driving EMT but that JNK was increasingly important for tumor cell EMT when MEK is substantially inhibited.

**Figure 7.**
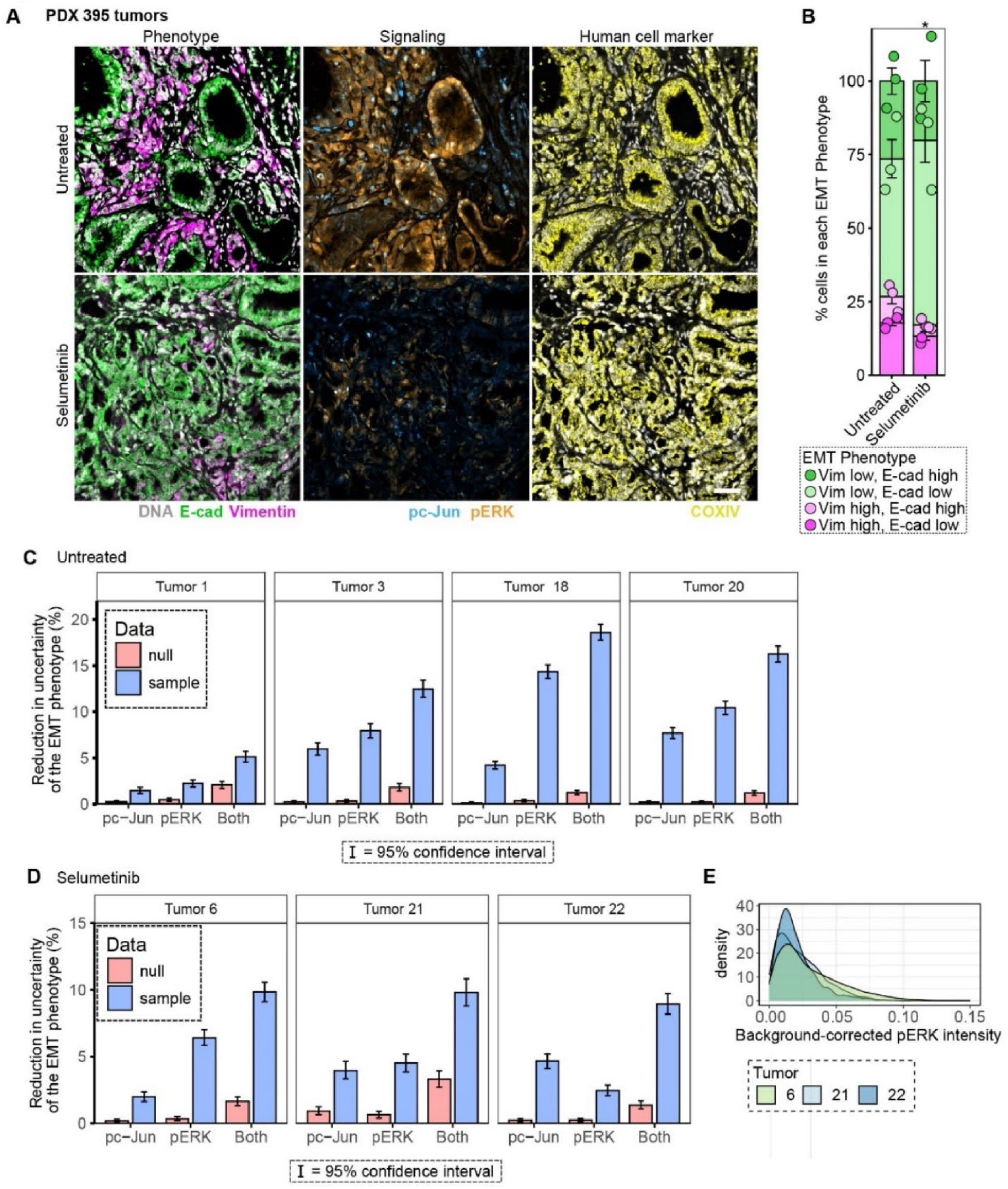
A mutual information model of EMT heterogeneity in patient-derived xenograft tumors supports the prominent role of ERK. **A)** Mice bearing PDX 395 tumors were untreated or treated with the MEK inhibitor selumetinib. Tumor sections were subjected to three rounds of 4i for the indicated proteins. Representative images are shown, n = 3. **B)** For the experiment described in (A), the mosaic plot displays percentages of cells in each EMT phenotype bin. n = 3, Chi-squared test, *p < 0.05, and error bars represent standard error of the mean. **C)** and **D)** For the control or selumetinib-treated mice described in (A), the percent reduction of uncertainty of the EMT phenotype from pERK, pc-Jun, or both signals from the ordered data or a randomly shuffled null dataset is plotted. n = 4 for untreated tumors, m=3 for selumetinib-treated tumors. Error bars show the 95% confidence interval of bootstrapped data. **E)** For the experiment described in (A), pERK intensity of background-subtracted selumetinib-treated tumors is plotted. Scale bar indicates 50 μm.

## DISCUSSION

We found that a network of signaling pathways was activated heterogeneously among pancreas cancer cells in response to growth factors or chemotherapeutics but that intrapopulation variation in the activation state of ERK specifically played a dominant role in explaining EMT heterogeneity. This experimentally validated model prediction contrasts starkly with a model prediction based on bulk signaling measurements that JNK signaling is the stronger determinant of growth factor-driven EMT. The discrepancy in predictions from models based on bulk versus single-cell measurements highlights the dangers of using population-level measurements to identify signaling pathways that control heterogeneous phenotypes. The errors that can arise may lead to improper drug target identification or incorrect conclusions about how druggable pathways work together. For example, it has been reported that ERK and JNK cooperate to drive EMT (*38, 40, 49*). We found that the ostensible cooperation between ERK and JNK in growth factor-mediated EMT that was apparently revealed through pharmacological inhibition studies arose due to JNK-mediated compensation in response to strong MEK inhibition. This important conclusion could not have been reached without single-cell measurements and the MI model. Similar compensation effects may be at play in the reported cooperation between ERK and JNK in regulating cell fate after DNA damage (*50*) and apoptosis (*51*).

EMT regulation by chemotherapy is far less well studied than in response to growth factors. We were surprised that ERK played a dominant role over pathways that are more generally associated with DNA damage stress responses, but other work that did not model or account for heterogeneities is at least qualitatively consistent with our conclusions. For example, in chemoresistant pancreas cancer cells that survived chronic gemcitabine exposure, MEK inhibition significantly reduced EMT and increased chemosensitivity(*26*). The question of how chemotherapy drives ERK activity remains open, but several avenues exist for chemotherapy-induced JNK activation via the DNA damage response, including suppressed expression of the phosphatase MKP-1 (*52*). One study posited that stochastic ERK activation controls cell cycle arrest and senescence in the context of DNA damage (*53*). Thus, ERK control of EMT in response to chemotherapy could arise simply through random pathway activation rather than via a robust pathway initiated by DNA damage.

The MI values in our models reached as high 30%, indicating that most of the uncertainty in the EMT phenotype was unexplained by the measured pathways. Yet, the MI models correctly identified signaling nodes that strongly regulated EMT. In prior studies, MI values of ∼20% were observed for information between signaling pathway nodes or ∼80% for information between a growth factor and transcription factor (*54–56*). The MI variables in our models spanned a larger biological range than considered in other models in that kinases or their substrates were measured along with a complex phenotype coordinated by multiple transcription factors. The large degree of biological separation may be one factor that explains the somewhat modest magnitude of MI achieved in our models. Interrelated issues of timing and the measurement approach we used may have also affected MI model quality. It is possible, for example, that protein measurements at earlier time points that captured quick transient effects would have yielded more informative variances across the experimental conditions. However, EMT phenotypes took more time to develop than the earliest transient signaling effects, and the fixed-cell immunofluorescence approach we employed requires that observations of signaling pathways and phenotypes be made simultaneously. In future work, live-cell reporters for signaling pathways and phenotypic markers could overcome this limitation. That approach may be confounded by reduced sensitivity of reporters compared to antibody-based measurements or practical limitations on the engineering of cells with numerous reporters (*57*). Even with somewhat modest MI values, the models developed here were remarkably useful for identifying phenotype-regulating signaling pathways that were validated across multiple cell backgrounds.

Thus, more sophisticated systems including live-cell reporters may be unnecessary for developing useful MI models.

An important prospective application of computational models of signaling in general is the rational design of combination therapies (*58*). Models based on population-level measurements of signaling can certainly reveal nonintuitive druggable circuits and processes in cancer cells [e.g., (*59*)], but models that directly discretize effects at the single-cell level are needed as phenotypes become increasingly heterogeneous (*60*). Our MI models of EMT heterogeneity pointed primarily to MEK inhibitors as EMT antagonists that will promote chemosensitivity, but a need to inhibit compensatory JNK signaling and the potential utility of SFK inhibition to target multiple MAPKs simultaneously were also identified. Pharmacological agents exist to test this concept, with four MEK inhibitors FDA-approved for solid tumors (trametinib, selumetinib, cobinetinib, and binimetinib) (*61*)and two SFK inhibitors approved for hematological malignancies (dasatinib and bosutinib) (*62*). Multiple JNK inhibitors have been tested in clinical trials. Of course, MEK inhibitors have already been explored in PDAC for their potential ability to synergize with inhibitors of autophagy, other kinases, or immune checkpoints (*63–65*). Preclinical studies using chemotherapy and trametinib together or sequentially have demonstrated cooperative tumor growth suppression and survival benefits (*66, 67*). However, a randomized controlled trial of trametinib and gemcitabine in patients with untreated metastatic pancreas cancer showed no significant differences in overall survival or progression-free survival (*68*).

When considering how to effectively combine MAPK inhibitors with chemotherapy to overcome EMT, dose scheduling and frequency will likely play important roles due to the possibility of multiple, potentially interfering, effects of MAPK pathway inhibition. MEK inhibition induces G1 cell cycle arrest (*69*), which limits gemcitabine incorporation into DNA (*70*). The effect of this was observed in a patient-derived xenograft mouse model of biliary cancer, where 48-hr selumetinib treatment followed by 48-hr washout for cell cycle release prior to gemcitabine treatment markedly improved tumor control compared to continuous co-treatment or gemcitabine alone (*71*). The effect of selumetinib on EMT markers was not assessed, but it seems clear that dosing schedules that allow for EMT antagonism while avoiding undesirable effects on cell cycle should be the objective. A related issue is the sensitivity of different cellular phenotypes to the degree of MAPK inhibition. At least 90% ERK inhibition is needed for effective antagonism of cell proliferation in some settings (*72*). Yet, we found that just a 30% population-level reduction in ERK activity could inhibit gemcitabine-induced EMT. Thus, even if a more continuous co-treatment approach were needed for EMT antagonism, there may be a window for low-dose trametinib to prevent chemotherapy-induced EMT without the undesirable consequence of slowing proliferation. Interestingly, while an ability for JNK to compensate for MEK inhibition was observed for growth factor-mediated and PDX tumor microenvironment-mediated EMT, JNK did not appear nearly as informative of EMT status for chemotherapy treatments of cells. Thus, careful optimization of the use of MEK inhibitors with chemotherapy, based on considerations discussed here, may be sufficient to improve patient outcomes.

## MATERIALS AND METHODS

### Cell culture

HPAF-II human pancreas cancer cells (Carl June, University of Pennsylvania) and human patient xenograft-derived PDX 395 cells (Todd Bauer, University of Virginia) were maintained in RPMI with 10% fetal bovine serum (FBS) (VWR), 1 mM L-glutamine (ThermoFisher), 100 units/mL penicillin (ThermoFisher), and 100 μg/mL streptomycin (ThermoFisher). KPCY-derived 7160c2 cells (Ben Stanger, University of Pennsylvania) were maintained in DMEM with 10% FBS, 1mM L-glutamine, 100 units/mL penicillin, and 100 μg/mL streptomycin. HPAF-II and PDX cell lines were tested for mycoplasma using the MycoAlert PLUS Detection Kit (Lonza) (2024, 2022). HPAF-II cells were authenticated via short tandem repeat profiling by the Genetic Resources Core Facility at Johns Hopkins University (2022). Recombinant human EGF, HGF, and TGFβ1 (Peprotech) were used at 50 ng/mL, 50 ng/mL, and 10 ng/mL, respectively. Gemcitabine (University of Virginia Clinical Pharmacy) and 5-fluorouracil (MP Bio) were used at 10 ng/mL and 10 μM, respectively. Wherever multiple conditions were used, growth factors, inhibitors, and/or chemotherapeutics were combined simultaneously, unless otherwise noted.

For treatments, complete medium containing growth factors, inhibitors, or chemotherapy drugs was replenished every 24 hr. The MEK inhibitor trametinib (Apex Bio) was used at 200 nM (HPAF-II), 20 nM (PDX 395), and 2 nM (7160c2) concentrations. The MEK inhibitor CI-1040 (LC Laboratories) was used from 0.001 μM to 1 μM. The JNK inhibitor SP600125 (LC Laboratories) was used at 10 μM, and JNK-IN-8 (MedChemExpress) was used at 5 μM. Stocks of all inhibitors were prepared in DMSO, and in experiments where inhibitors are used, total DMSO concentrations were brought up to 0.1% across conditions. Cell seeding densities in 96-well plates were chosen to enable sufficient cell number for analysis while maintaining sub-confluence at the endpoint across different cell lines and treatments/timings. For example, 4,000 HPAF-II cells were seeded per well for experiments involving 5-day growth factor treatments, but 12,000 HPAF-II cells were seeded per well for 2-day maximum treatments. When multiple time points were sampled on the same 96-well plate, all treatments were scheduled to conclude simultaneously. For example, for a comparison of 1-5 days of growth factor treatment on a 96-well plate, cells were plated simultaneously, and the 5-day treatment was begun first, followed by the 4-day treatment the next day, and so forth.

### Antibodies

Antibodies against E-cadherin (Invitrogen 13-1900), vimentin (Santa Cruz sc-373717), ERK (CST 4695), pc-Jun Ser73 (CST 3270), and pERK1/2 Thr202/Tyr204 (CST 4370) were used for immunofluorescence and western blotting. For immunofluorescence, antibodies against pp38 Thr180/Tyr182 (CST 4511), pSrc Tyr419 (Santa Cruz sc-81521), SMAD2/3 (Santa Cruz sc-133098 AF 546), STAT3 (CST 9139), and NFκB (p65 regulatory subunit, CST 8242) were used. For western blotting, antibodies against GAPDH (Santa Cruz sc-32233) and GRB2 (Santa Cruz sc-255) were used. Antibody dilutions for western blotting were used at manufacturer recommendations. For PDX tumors, antibodies against E-cadherin (Invitrogen 13-1900), vimentin (Santa Cruz sc-373717), pc-Jun Ser73 (CST 3270), and pERK Thr202/Tyr204 (CST 4370) were used as well as conjugated anti-COXIV (3E11)-Alexa Fluor 594 (CST 8692).

### Western blots

Cells plated in 6-well plates were lysed using a cell extraction buffer (ThermoFisher, FNN0011) supplemented with protease and phosphatase inhibitors (Sigma-Aldrich; P8340, P5726, and P0044). Crude lysates were centrifuged at 20,000×g for 10 min at 4°C. Supernatants were collected as clarified lysates, and total protein concentrations were determined with a micro-bicinchoninic acid (BCA) assay (Thermo Fisher, 23225). Clarified lysate, 10× DTT, 4x LDS sample buffer (ThermoFisher, NP0007), and MilliQ water were combined to ensure equal protein amounts and equal sample volumes, and samples were then heated for 10 min at 100°C. For electrophoresis, samples were loaded on 1.5-mm NuPAGE gradient (4-12%) gels (Invitrogen, NP0336). After electrophoresis, gels were transferred to a 0.2-μm nitrocellulose membrane using the TransBlot Turbo Transfer System (BioRad). Membranes were blocked at room temperature for 1 hr with shaking, washed, incubated with primary antibodies diluted in intercept blocking buffer (IBB, Licor, 927-60001) overnight at 4°C, washed, and then incubated with secondary antibodies diluted in IBB at room temperature for 2 hr. All wash steps were performed as three times for 5 min each with 0.1% Tween-20 in PBS and gentle shaking. Membrane imaging was performed on a LI-COR Odyssey CLx. Membrane stripping was performed with 0.2 M NaOH as needed and confirmed via reimaging. GAPDH intensity or GRB2 and ERK average intensities were used as loading controls.

### siRNA-mediated knockdowns

Cells were reverse-transfected with siRNA according to manufacturer protocol using Lipofectamine RNAiMAX (ThermoFisher). Cells were plated on top of the siRNA-lipid complexes 24 hr prior to treatment and fixed for immunofluorescence staining 72 hr post-transfection. siRNA against ERK1/2 (#6560) was purchased from Cell Signal Technology. c-Jun (sc-29223) and control siRNA (sc-37007) were purchased from Santa Cruz Biotechnology.

### Iterative immunofluorescence staining

The original 4i protocol (*36*) was followed except for the use of IBB in the blocking and antibody staining steps. The published protocol contains recipes for all buffers described here. 96-well plate washes were performed using a BioTek EL406 washer dispenser, washing 6× per wash step by removal and replacement of the top 160 μL, leaving 40 μL volume as a mechanical buffer to prevent cell loss due to fluid shear. Cells were fixed in 4% paraformaldehyde in PBS, permeabilized in 0.5% Triton-X-100 for 15 min and washed in deionized (DI) water. The following steps were then performed in order for each round of imaging: incubation with elution buffer for 3× 10 min, washing with 1× PBS, blocking in IBB + 150 mM maleimide for 30 min at room temperature, staining with primary antibody in IBB overnight at 4°C, washing with 1× PBS, staining with secondary antibody in IBB 1 hr at room temperature, washing with DI water, imaging in imaging buffer, and then washing with DI water prior to the next round of antibody elution. The 6-signal and 10-signal antibody panels were split across rounds of 4i **(SuppTables S1 and S2)**.

For tumor sections, 4i was performed as described above but with several modifications. Elution was performed using 3× 5-min washes, and wash steps were conducted using 3× 5-min washes through gentle manual pipetting. Samples were imaged using imaging buffer + 10% glycerol. Coverslips were placed on the slides just before imaging, and slides were immersed in deionized (DI) water just after imaging to gently release the coverslip from the tumor sample. Antibody dilutions for tissue 4i are described in **SuppTable S3**.

### Immunofluorescence staining

Non-iterative immunofluorescence was performed by washing cells 2× in PBS, fixing in 4% paraformaldehyde for 20 min, then washing 3× in PBS before permeabilization with 0.25% Triton-X-100 in PBS for 5 min. Cells were then washed 3× in PBS, and incubated overnight at 4°C with primary antibodies diluted in IBB using the same dilutions as in 4i staining (Supplementary Tables S1, S2). Cells were then washed 3× in PBS, then stained with secondary antibodies diluted in IBB for 1 hr at room temperature. Cells were then washed 3× in PBS and then imaged in 20% glycerol in PBS.

### Live/Dead staining

Experiments for live/dead staining used either TO-PRO-3 (Thermo Fisher, T3605) or Sytox Blue (Thermo Fisher, S34857) for dead cell staining and phase contrast imaging or SiR-DNA (Cytoskeleton, CY-SC007) to identify all cells. Each stain was used at a 1:1000 dilution in PBS. Cells were first washed once with PBS before staining in the dark at 37°C for 30 min. Cells were washed twice in PBS before imaging in complete medium without phenol red.

### Fluorescence microscopy and automated image analysis for cell culture

Cells in 96-well plates were imaged using a BioTek Cytation5 plate reader in “Experiment-mode” for high-throughput automated imaging. Images were captured using a 10× phase contrast objective, and Gen5 imaging software was used to produce tiff files of individual fluorescence channels. Imaging protocols were programmed to capture two rows of three frames spaced evenly apart in each well, using auto-focus to find the appropriate focal planes across _the_ plate. Images with debris or imaging artifacts were discarded before image analysis. At least 1000 cells were quantified for each well.

Illumination correction and background subtraction were performed using the Cytation5 automated rolling ball background subtraction data reduction protocol. CellProfiler v4.2.1 (Broad Institute) was used for image alignment, cell segmentation, and quantification of signal intensity and localization (*73*). To correct for camera drift across rounds of 4i, images were first aligned using the CellProfiler *align* module. Images of the DNA stain Hoechst (DAPI channel) for each of the imaging rounds were aligned separately, cropping nonoverlapping regions to create a reference image of fully overlapping nuclei. For each round of images, the original DNA stain image was then aligned separately to the reference image with the other channels in that round being aligned similarly as the DNA image based on the degree of *x*/*y* translation needed to align the DNA image. For cell segmentation, cells were first identified using the DNA stain Hoechst (Identify Primary Objects), and cell boundaries were then identified using total ERK (Identify Secondary Objects). Intensity measurements were integrated over whole cells or nuclei. Where plotted, mean signal intensities for IF images indicate the mean of the integrated intensities of all cells within a replicate.

### Fluorescence microscopy and automated image analysis for tumor sections

After antigen retrieval and antibody staining, tumor sections on charged glass slides were imaged using a Zeiss LSM 880 confocal microscope in the UVA Advanced Microscopy Facility. Images were taken using a Plan-Apochromat 20×/0.8 M27 objective, and ZEN Black imaging software was used to produce .czi files. At least eight frames per tumor section were taken. Images were acquired as *z*-stacks across the thickness of the tumor in 1-μm intervals, then rendered as 2D images through maximum intensity projection in Fiji (*74*). Segmentation of nuclei and cells was performed using the deep learning-enabled segmentation algorithm *Mesmer* through the DeepCellKiosk Fiji plugin (*75, 76*). The first output of cell-occupied, shaded regions was saved as an image. The segmentation lines were added to the blank image as an overlay, then flattened and converted into a binary mask and saved as a separate image. These images were included in CellProfiler, where shaded cellular regions were thresholded into binary images. Cell segmentation masks were then subtracted from the cell shaded regions, resulting in clear cell segmentation easily interpretable by the *Identify Primary Objects* module.

These objects were then used to measure object intensities for all other channels. Intensity measurements were made as mean pixel intensities across whole cells or nuclei. To compare signal intensities across tumor sections, a value equal to the lowest 10 percent of signal intensities measured across each image was subtracted from the intensity measurements from each cell. This provided a way of accounting for inter-tumoral staining variations.

### Patient-derived xenograft tumors

Tumor xenograft experiments performed using the MEK inhibitor selumetinib were described by Brown et al.(*40*) Briefly, mice bearing tumors derived from PDX 395 were treated with selumetinib (2.5 mg/kg, orally, twice daily; LC Laboratories) or vehicle control (0.5% hydroxypropylmethylcellulose + 0.1% Tween 80 in water). Excised tumors were fixed in 10% zinc-buffered formalin for 24 hr, transferred to 70% ethanol, and then embedded in paraffin. Paraffin-embedded tumors were cut into 4-5 μm sections onto charged slides. The UVA Research Histology Core performed tumor embedding and sectioning. For antigen retrieval, slides were immersed in xylene for 2× 10 min at room temperature, then transferred to 100%, 90%, and then 80% ethanol each for 5 min at room temperature. Slides were then rinsed in ultrapure water and antigen-retrieved with high pH at 95°C for 20 min.

### Gaussian mixture modeling

Gaussian mixture modeling was performed using the *ClusterR* package to determine the optimal number of cell signaling clusters (*77*). For the dataset in Figure 1, pERK and pc-Jun intensities were each fitted to gaussian mixture models with up to 10 clusters. The optimal number of clusters was chosen by examining the point at which the Bayesian Information Criterion was not substantially lowered through further binning.

### Binning of signals from fluorescence microscopy

Phenotype bins for vimentin, TO-PRO-3, and Sytox blue were created by measuring the median signal intensity for the untreated condition and adding twice the median absolute deviation of the signal in the untreated condition to create a cutoff, above which the phenotypes were classified as “positive” or “high.” For E-cadherin, which was expressed well at baseline in the cells we analyzed, we chose a cutoff that best captured the separation of membranous E-cadherin intensities between untreated cells and day-5 growth factor-treated cells: median E-cadherin membrane intensity minus the median absolute deviation for the untreated condition. Above that cutoff, E-cadherin was classified as “high.”

Signal intensities (for phosphosignals) or signal nuclear-to-cytoplasmic ratios (for total proteins) were binned into four groups based on quartile cutoffs for the range of activation states over the full experiment: high, medium-high, medium-low, or low. Four was chosen as the number of groups based on Gaussian mixture modeling (Supplementary Figure S3).

### MI model development

MI calculates the reduction of uncertainty (or entropy) of one variable that occurs from knowing another variable within a population. For example, if two variables always exhibit the same state or value across observations (e.g., cells) in a population, the MI between them is 100%. Uncertainty (*H*) is a unitless quantity based on outcome probabilities. A variable *X* can have *N* states *x_i_*, where the index *i* ranges from 1 to *N*, and *x_i_* represents a discrete possible state value for *X* (e.g., top 25% of the range of observable intensities for *X*). With just one variable *X*, the uncertainty of that variable *H*(*X*) is calculated using the probability distribution of the occurrence for each state *p*(*x_i_*) as:

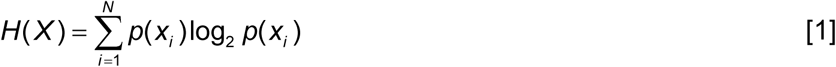

which is zero for a single possible state for *X* (i.e., *N* = 1, *p*(*x_i_*)= 1). For two variables *X*_1_ and *X*_2_, the relevant probability distribution is *p*(*x_i_,*x*_j_*), which captures the likelihoods of joint occurrences for each state *x_i_* of *X*_1_ and each state *x_j_* of *X*_2_. The joint uncertainty of *X*_1_ and *X*_2_, *H*(*X_1_*,*X_2_*), can be calculated as:

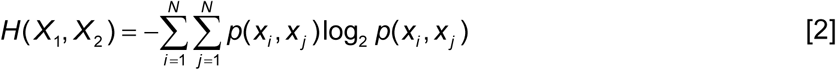

In Eq. [2], we have assumed the same number of possible states for *X*_1_ and *X*_2_, but the variables assessed in MI calculations can in general have different numbers of states. We proceed with the assumption that the number of states for each variable is conserved. Because the uncertainty of a two-variable system is defined by *H*(*X*_1_,*X*_2_), the difference between *H*(*X*_1_,*X*_2_) and *H*(*X*_2_) is just the uncertainty of *X*_1_ given *X*_2_, *H*(*X*_1_│*X*_2_) (*78*):

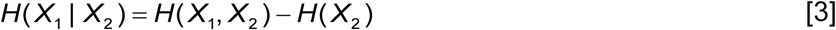

For an arbitrary number of variables *n*, the joint uncertainty can be calculated as:

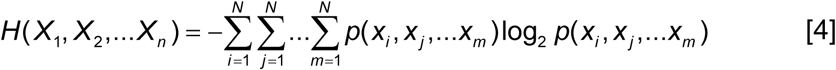

where the index *m* is used for the states *x_m_* of the n^th^ variable *X_n_*. For *n* variables then, the uncertainty of *X*_1_ given all other variables is:

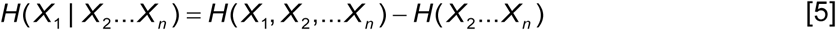

The percent reduction in uncertainty of knowing the state of *X*_1_ achieved by knowing the states of *X*_2_…*X*_n_ is given by:

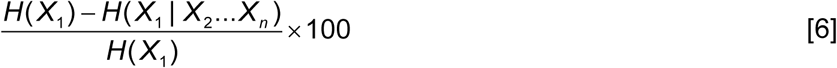

where *H*(*X*_1_) can be computed using Eq. [1].

This general development can be applied to analyze the regulation of EMT by multivariate signaling by treating *X*_1_ as an EMT variable and *X*_2_…*X_n_* as signaling variables. In our application of that approach for the calculations we performed, *N* = 4 for all variables. As described in *Methods*, the four EMT states were defined based on measurements of vimentin and membranous E-cadherin. For each signaling node, the four states were defined based on quartiles of the entire range of signals measured for that signal across all conditions.

To create null datasets for comparisons, data were shuffled using the *sample_n()* function in R. Both the real data and the null data were bootstrapped by randomly resampling 80% of the data 1000 times (replacing the data after it was chosen to avoid biasing the output) to obtain 95% confidence intervals. All calculations were performed using code developed in the R programming language.

### Cell population-level modeling

Western blot intensity measurements for each of the signaling and phenotype markers were divided by their respective loading controls (GAPDH for pc-Jun, total ERK for pERK, or averaged intensities of two loading controls, ERK and GRB2, for E-cadherin and vimentin) and analyzed in one of two ways.

To measure activation of signaling pathways and change in expression of EMT markers at the population level, data was assessed on the same scale across separate western blots for different and biological replicates by subtracting the median value of the replicate from each data point within the replicate. To then ensure only non-negative values were being compared, the minimum value was subtracted from each data point.

To perform the multiple linear regression, loading control-normalized western blot intensity measurements were mean-centered and variance-scaled. The EMT variable was generated by subtracting the rescaled E-cadherin measurements from the rescaled vimentin measurements, which also served as the relative EMT score. The model was generated using the *lm()* function in base R assuming a model form of EMT = β_0_ + β_1_[pc-Jun] + β_2_[pERK], where β_0_ is the *y*-intercept and β_1_ and β_2_ are the slope coefficients (partial regression coefficients) for independent variables [pc-Jun] and [pERK]. Added-variable plots to visualize the partial regressions were constructed using the *avPlots* function from the *car* package. Diagnostic plots were created using the *plots()* function from the R stats package on the linear model.

### Statistical analysis

The statistics package in base R was used. Samples were compared using ANOVA with Tukey’s honestly significant difference. For conditions where only one comparison was needed, unpaired t-tests were used. For statistics on categorical variable distributions, the Chi Squared Test was used. ggplot2 was used to create figures. Statistical analyses were performed using R version 4.3.0.

## ACKNOWLEDGEMENTS

We thank Dr. Mohammad Fallahi-Sichani for assistance with learning the 4i protocol and Dr. Ben Stanger for providing the KPC-derived cell line. The authors acknowledge the resources and services provided by Research Histology Core Facility, a shared facility in the School of Medicine at the University of Virginia. This work also used the Zeiss LSM 880 in the Advanced Microscopy Facility which is supported by the University of Virginia School of Medicine, Research Resource Identifiers (RRID): SCR_018736. Figure 1A was created using BioRender.com.

## AUTHOR CONTRIBUTIONS

**M.C.B.**: Conceptualization, funding acquisition, investigation, methodology, data analysis, writing-original draft, and editing. **B.A.B.** Investigation and methodology. **S.J.A.**: Investigation, methodology, writing-review and editing. **T.W.B.**: Resources, supervision, writing-review and editing. **M.J.L.**: Conceptualization, supervision, funding-acquisition, project administration, writing-original draft, writing-review and editing.

## DATA AND MATERIALS AVAILABILITY

All analysis for this paper was conducted using code written in R programming language. All code generated will be available on the Lazzara lab github site.

## SUPPLEMENTAL MATERIALS

### Supplemental Methods

RNA from HPAF-II and 7160c2 cells was extracted using the RNeasy Kit (Qiagen) and reverse-transcribed using HighCapacity cDNA Reverse Transcription Kit (Applied Biosciences). qRT-PCR was performed using PowerUp SYBR Green (Applied Biosciences) per manufacturer protocol using a QuantStudio3 system (Applied Biosystems). qRT-PCR primer sequences are provided in **SuppTable S4**. Measurements were analyzed with the ddCt method. Gene expression data are displayed as normalized fold-changes, using *GAPDH* for normalization.

**Supplemental Table S1:** 6-signal 4i panel.

| Antibody target | Species | Dilution | Secondary | Round in 4i |
| --- | --- | --- | --- | --- |
| Hoechst | N/A | 1:2000 | N/A | All |
| pc-Jun | rabbit | 1:400 | α-rabbit 647 | 1 |
| pERK | rabbit | 1:200 | α-rabbit 647 | 2 |
| ERK1/2 | rabbit | 1:800 | α-rabbit 546 | 3 |
| E-cadherin | rat | 1:800 | α-rat 488 | |
| Vimentin | mouse | 1:800 | α-mouse 647 | |

**Supplemental Table S2:** 10-signal 4i panel.

| Antibody target | Species | Dilution | Secondary | Round in 4i |
| --- | --- | --- | --- | --- |
| Hoechst | N/A | 1:2000 | N/A | All |
| pSrc | mouse | 1:200 | α-mouse 546 | 1 |
| pp38 | rabbit | 1:1600 | α-rabbit 647 | |
| Smad2/3 | mouse | 1:800 | 546 conjugate | 2 |
| pc-Jun | rabbit | 1:400 | α-rabbit 647 | |
| Stat3 | mouse | 1:800 | α-mouse 546 | 3 |
| pERK | rabbit | 1:200 | α-rabbit 647 | |
| NFkB | rabbit | 1:400 | α-rabbit 546 | 4 |
| E-cadherin | rat | 1:800 | α-rat 488 | |
| Vimentin | mouse | 1:800 | α-mouse 647 | |

**Supplemental Table S3:** PDX 395 tumor section 4i panel.

| Antibody target | Species | Dilution | Secondary | Round in 4i |
| --- | --- | --- | --- | --- |
| Hoechst | N/A | 1:1000 | N/A | All |
| pc-Jun | rabbit | 1:200 | α-rabbit 647 | 1 |
| E-cadherin | rat | 1:100 | α-rat 546 | |
| pERK | rabbit | 1:200 | α-rabbit 647 | 2 |
| Vimentin | mouse | 1:200 | α-mouse 647 | |
| COXIV | rabbit | 1:200 | α-rabbit 647 | 3 |

**Supplemental Table S4:**
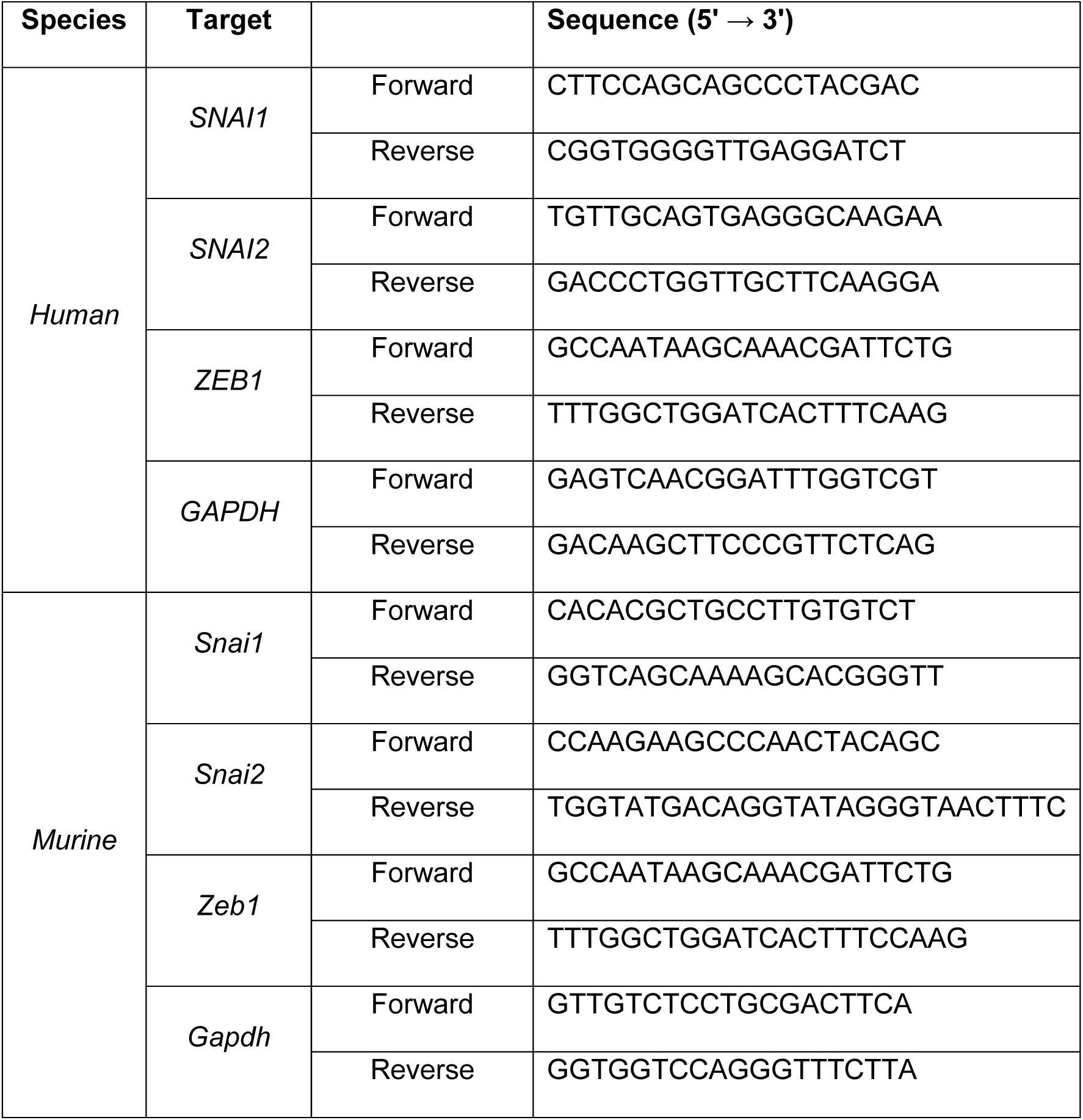
qRT-PCR primers.

### SUPPLEMENTAL FIGURE LEGENDS

**Supplemental Figure S1.**
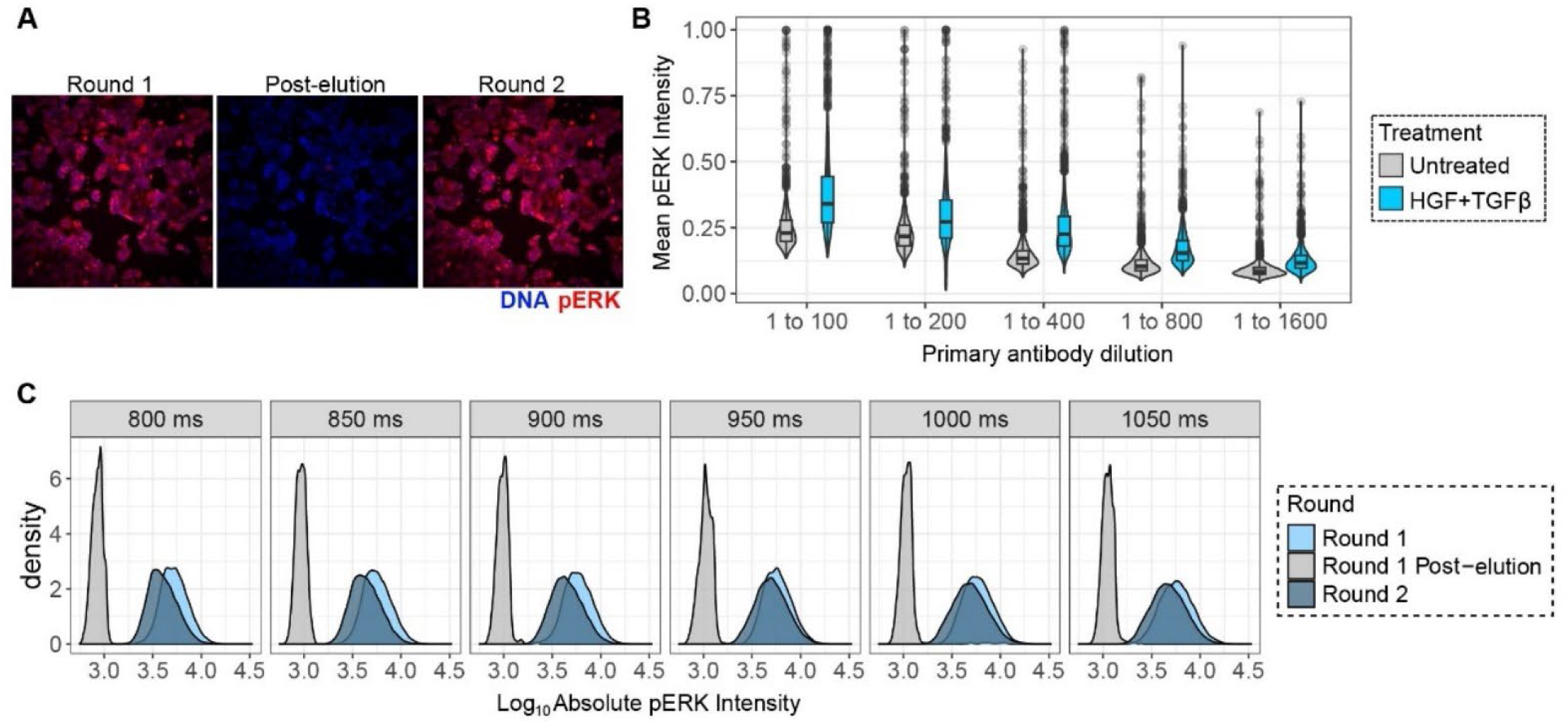
Antibody optimization steps for 4i. **A)** Representative images of pERK signal in HPAF-II cells (completed medium, untreated) over two rounds of 4i. Image representative of n = 3. **B)** Violin and box plots of pERK intensity in HPAF-II cells treated with or without 50 ng/mL HGF + 10 ng/mL TGFβ for 30 min for the indicated primary antibody dilutions for round 1 of 4i staining, n = 3. **C)** Density plots displaying distribution of baseline pERK signaling intensities in untreated HPAF-II cells during round 1, round 1 post-elution, and round 2 of the 4i protocol over a range of imaging exposure times, n = 3. Scale bar indicates 300 μm.

**Supplemental Figure S2.**
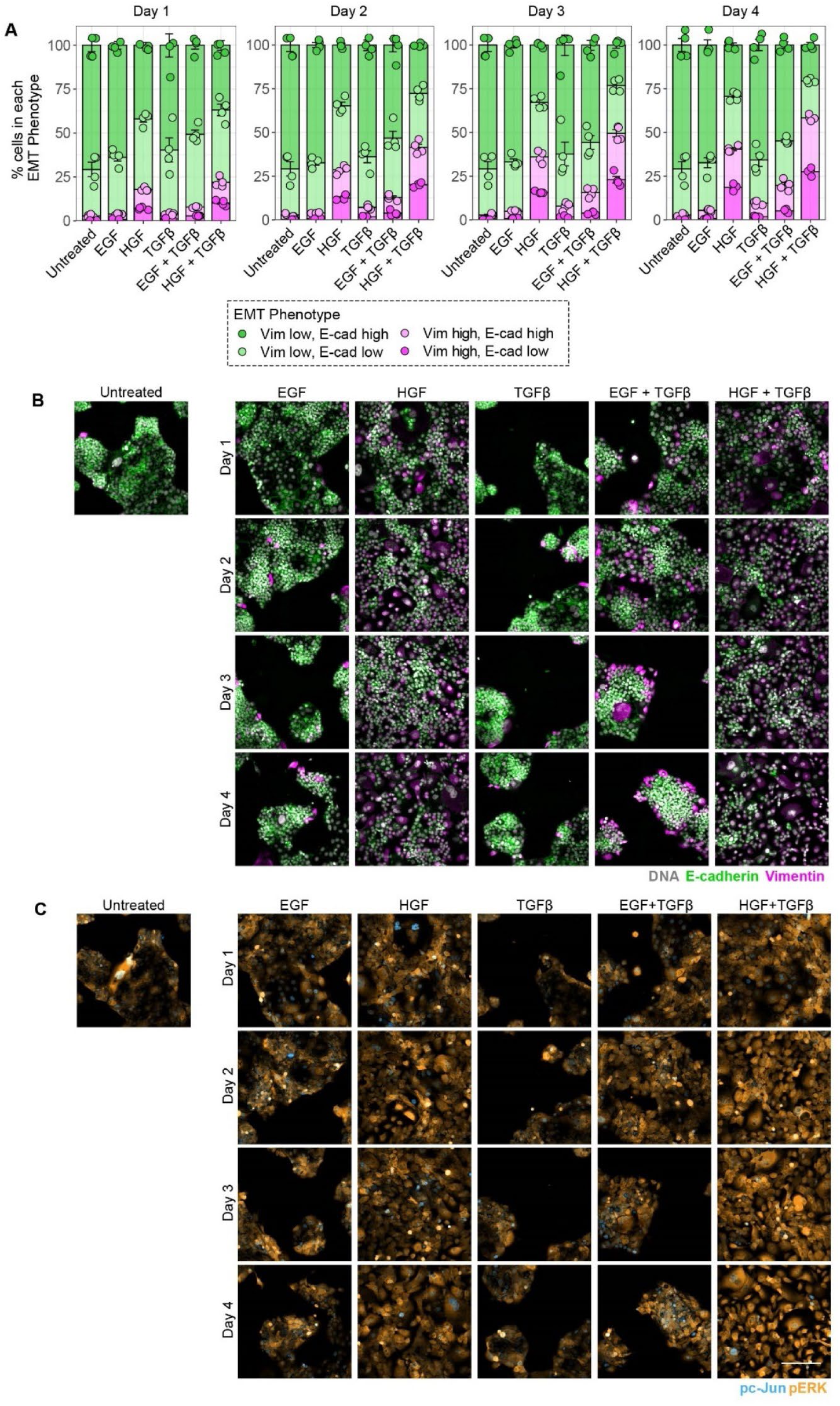
EMT induction for 1-4 days. HPAF-II cells were treated with 10 ng/mL TGFβ, 50 ng/mL EGF, 50 ng/mL HGF, EGF+TGFβ, or HGF+TGFβ for 1-4 days. **A)** Mosaic plot displaying percentage of cells in each EMT phenotype bin over 1-4 days, n = 4. **B)** Representative 4i images of vimentin and E-cadherin over 1-4 days. **C)** Representative 4i images of pERK and pc-Jun for 1-4 days. For all panels, error bars represent standard error of the mean. Scale bar indicates 300 μm.

**Supplemental Figure S3.**
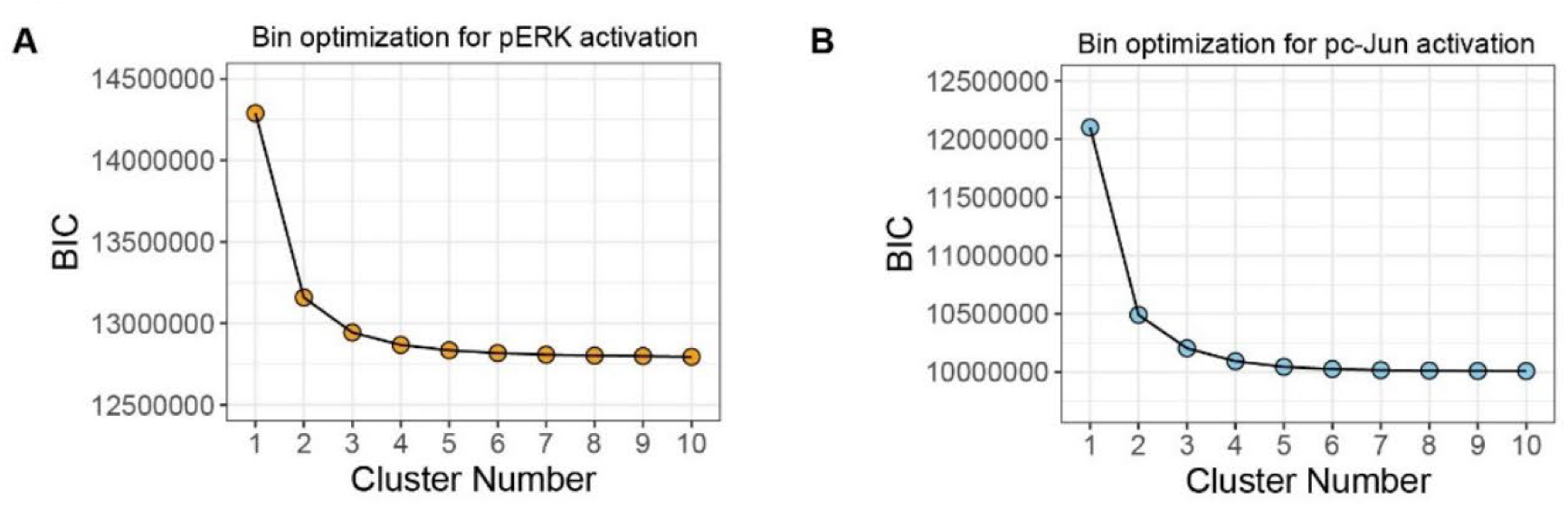
Cluster number identification from Gaussian mixture models of signaling data. HPAF-II cells were treated as described in Figure 1 for 1-5 days, n = 4. Gaussian mixture models were made for the pERK **(A)** and pc-Jun **(B)** signals. Line graphs display Bayesian information criterion (BIC) values versus cluster numbers for up to 10 clusters.

**Supplemental Figure S4.**
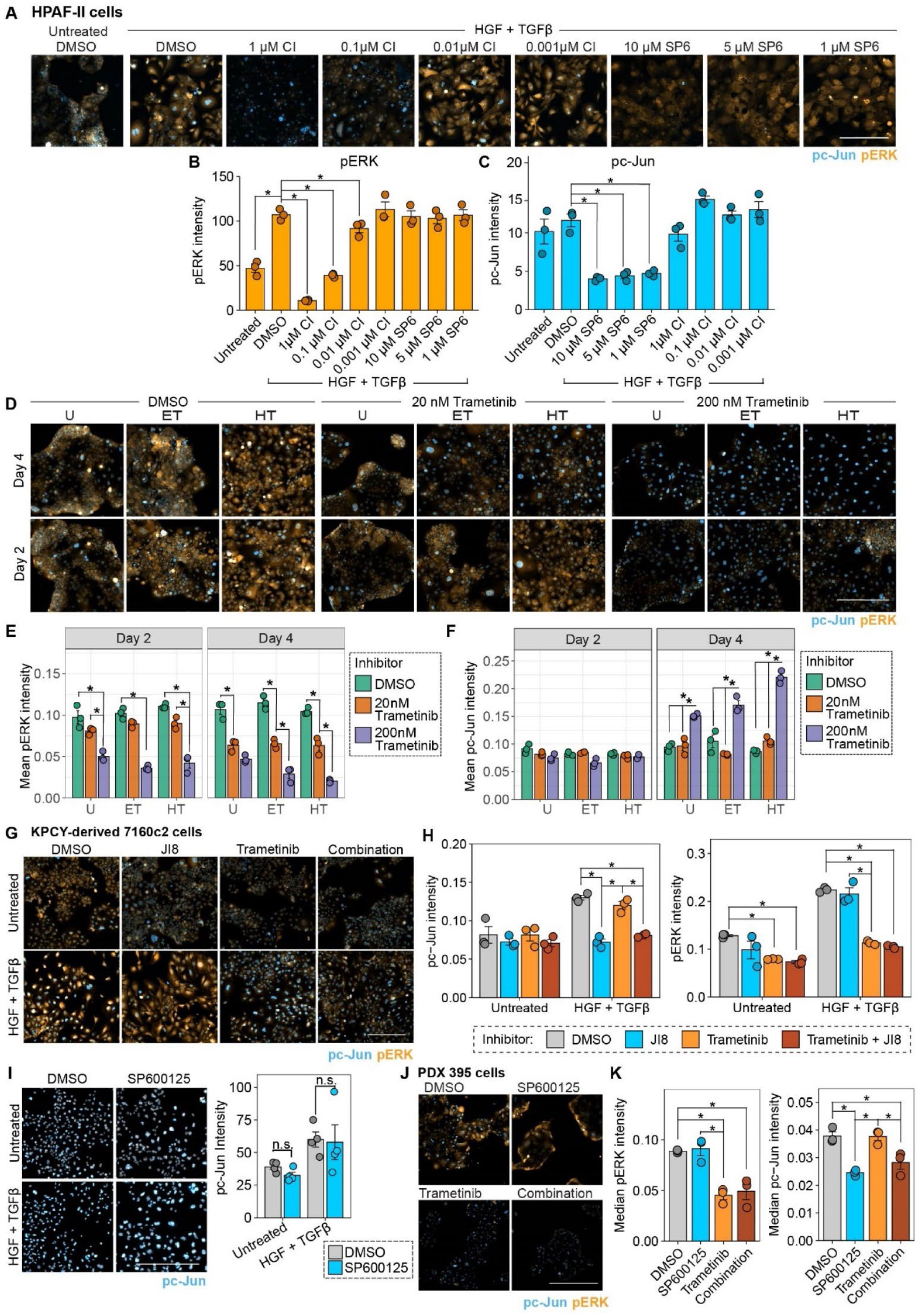
Selected inhibitor concentrations antagonize targets and minimize off-target effects. **A)** HPAF-II cells were treated with or without 50 ng/mL HGF + 10 ng/mL TGFβ, and HGF+TGFβ-treated cells were co-treated with DMSO or the indicated concentrations of the MEKi CI-1040 (CI) or the JNK inhibitor SP600125 (SP6) for 4 days. Representative 4i images of the indicated proteins are shown, n = 3. **B)** For the experiment described in (A), mean of pERK integrated intensity of cells is plotted. n = 3, one-way ANOVA with Tukey’s multiple comparisons test, comparisons shown against HGF+TGFβ with DMSO. **C)** For the experiment described in (A), pc-Jun intensity across cells is plotted. n = 3, one-way ANOVA with Tukey’s multiple comparisons test, comparisons shown against HGF+TGFβ with DMSO. **D)** HPAF-II cells that were treated with medium (U, untreated), 50 ng/mL EGF + 10 ng/mL TGFβ (ET), or 50 ng/mL HGF + 10 ng/mL TGFβ (HT) were treated with DMSO, 20 nM trametinib, or 200 nM trametinib for 2 or 4 days, as indicated. Representative 4i images for the indicated proteins are shown, n = 3. **E)** and **F)** For the experiment described in (D), pERK and pc-Jun intensity are plotted. n = 3, one-way ANOVA with Tukey’s multiple comparisons test. **G)** KPCY-derived 7160c2 cells that were treated with or without 50 ng/mL HGF + 10 ng/mL TGFβ were treated with DMSO, 5 μM JNK-IN-8, or 2 nM trametinib for 3 days. Representative 4i images for the indicated proteins are shown, n = 3. **H)** For the experiment described in (G), pc-Jun intensity (left) and pERK intensity (right) are plotted. n=3, one-way ANOVA with Tukey’s multiple comparisons test. **I)** KPCY-derived 7160c2 cells that were treated for 3 days with or without 50 ng/mL HGF + 10 ng/mL TGFβ were treated with DMSO or 10 μM SP600125. Representative 4i images of the indicated proteins are shown (left), and pc-Jun intensity is plotted (right). n = 4, one-way ANOVA with Tukey’s multiple comparisons test. **J)** PDX 395 cells were treated with DMSO, 10 μM SP600125, 20 nM trametinib, or a combination of both for 3 days. Representative 4i images of the indicated proteins are shown, n = 3. **K)** For the experiment described in (J), pERK intensity (left) and pc-Jun intensity (right) are plotted, n = 3, one-way ANOVA with Tukey’s multiple comparisons test. For all panels, *p < 0.05, and bars represent standard error of the mean. Scale bars indicate 300 μm.

**Supplemental Figure S5.**
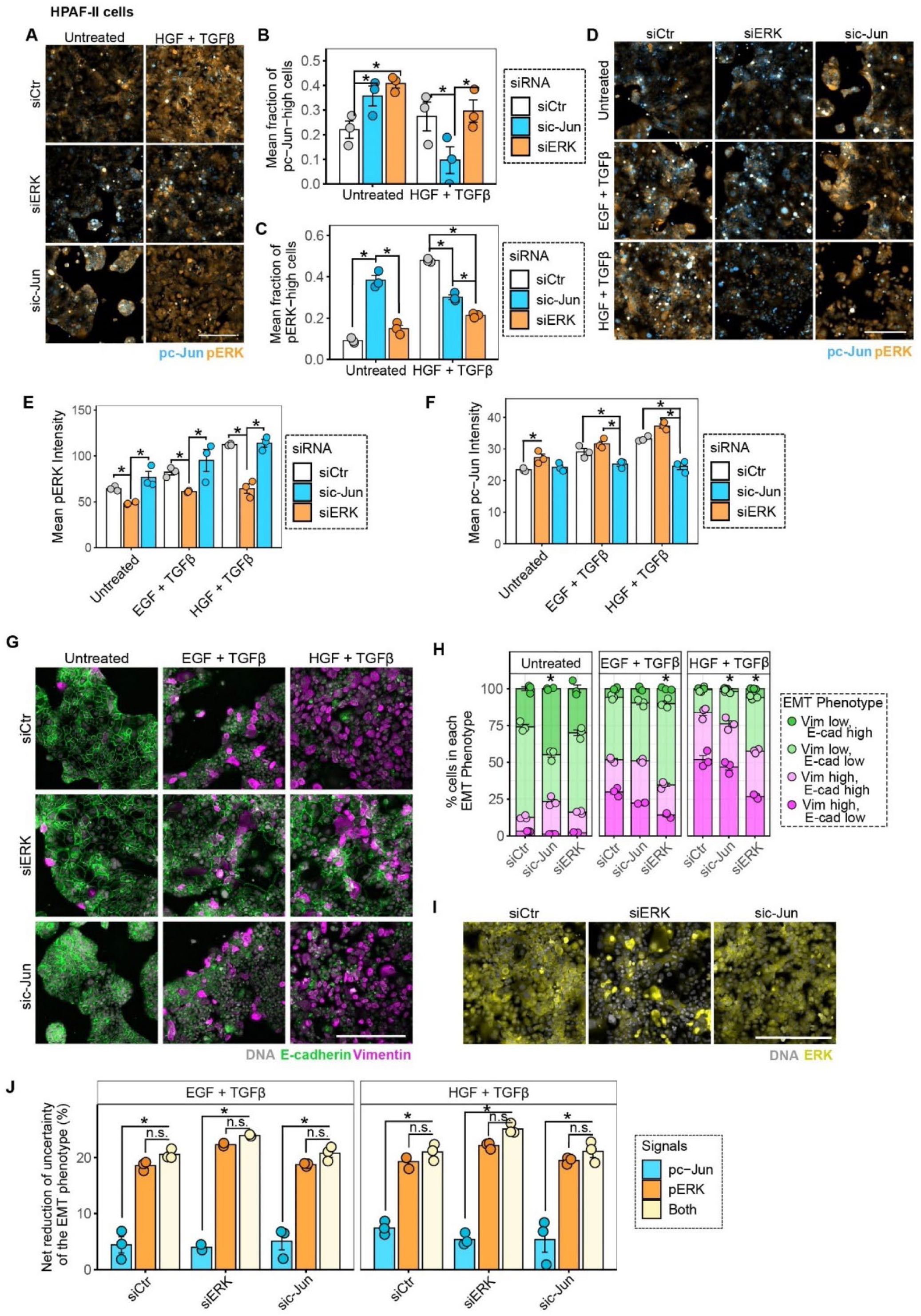
siRNA-mediated knockdown of ERK or c-Jun supports inferences from inhibition studies that ERK plays a dominant role over JNK in regulating EMT heterogeneity. HPAF-II cells were transfected with control siRNA or siRNAs targeting ERK1/2 or c-Jun for 24 hr and then treated with 50 ng/mL HGF + 10ng/mL TGFβ for 48 hr. **A**) Representative 4i images of pERK and pc-Jun for the indicated conditions. Images representative of n = 3. **B)** and **C)** Bar plots displaying fraction of cells in “pc-Jun high” or “pERK high” signaling bins for the indicated conditions. n = 3, one-way ANOVA with Tukey’s multiple comparisons test. **D)** HPAF-II cells were transfected with control siRNA or siRNAs for the knockdown of ERK1/2 or c-Jun for 24 hr and then treated with 50 ng/mL EGF + 10 ng/mL TGFβ or 50 ng/mL HGF + 10 ng/mL TGFβ for 24 hr. Representative 4i images of pERK and pc-Jun are shown, n = 3. **E)** and **F)** For the experiment described in (D), pERK or pc-Jun signal intensity is displayed. n = 3, one-way ANOVA with Tukey’s multiple comparisons test. **G)** HPAF-II cells were transfected with control siRNA or siRNAs for the knockdown of ERK1/2 or c-Jun for 24 hr and then treated with 50 ng/mL EGF + 10 ng/mL TGFβ or 50 ng/mL HGF + 10 ng/mL TGFβ for 48 hr. Representative 4i images of indicated proteins across treatment and siRNA conditions at 48 hr, n = 3. **H)** For the experiment described in (G), the mosaic plot displays percentages of cells in each EMT phenotype bin. n = 3, Chi-squared test. **I)** For the experiment described in (G), representative 4i images of total ERK are shown, n = 3**. J)** For the experiment described in (G), the net percent reduction in uncertainty of the EMT phenotype based on pERK, pc-Jun, or both signals is plotted. n=3, one-way ANOVA with Tukey’s multiple comparisons test. For all panels *p < 0.05, and error bars represent standard error of the mean. Scale bars indicate 300 μm.

**Supplemental Figure S6.**
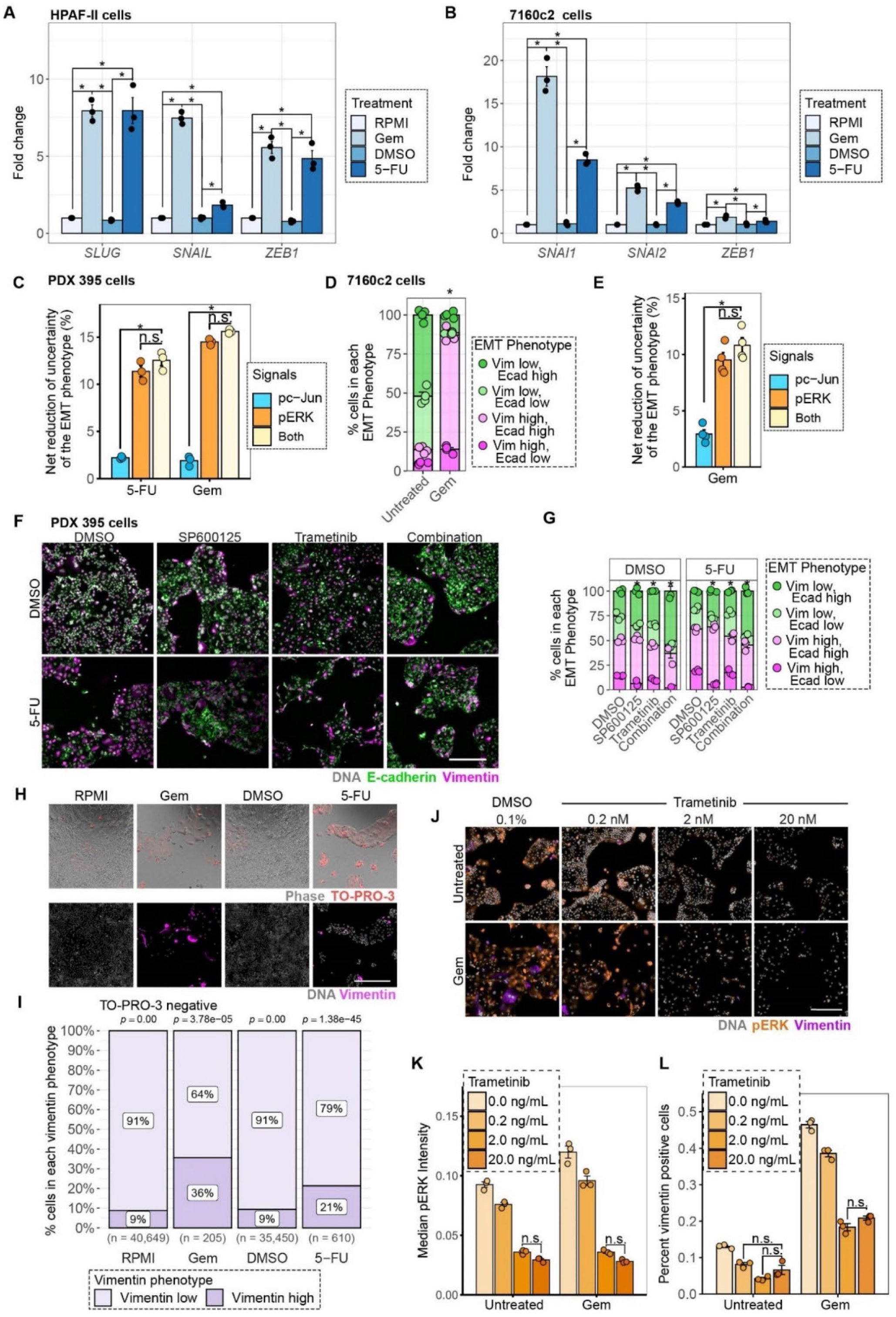
Chemotherapy drives a *bona fide*, persistent EMT. **A)** HPAF-II cells were treated with 10 ng/mL gemcitabine (Gem) or 10 ng/mL 5-fluorouracil (5-FU) for 48 hr. RPMI and DMSO were used as controls for gemcitabine or 5-FU, respectively. qRT-PCR was performed for the indicated EMT markers, and *GAPDH* was used for normalization. n = 3, one-way ANOVA test per transcript with Tukey’s multiple comparisons test. **B)** 7160c2 cells treated as described in panel (A) for 24 hr followed by media replacement for another 24 hr. qRT-PCR was performed for the indicated transcripts, with *GAPDH* used for normalization. n = 3, one-way ANOVA test per transcript with Tukey’s multiple comparisons test. **C)** PDX 395 cells were treated with 10 ng/mL gemcitabine or 10 ng/mL 5-FU for 3 days. The net percent reduction in uncertainty of the EMT phenotype based on pERK, pc-Jun, or both signals is plotted. n = 3, one-way ANOVA with Tukey’s multiple comparisons test. **D**) KPCY-derived 7160c2 cells were pulsed with or without 10 ng/mL gemcitabine for 24 hr followed by a complete media washout for 24 hr, and immunofluorescence microscopy was performed for EMT markers at 48 hr (shown in Figure 4F). The mosaic plot displays percentages of cells in each EMT phenotype bin. n = 4, Chi-squared test. **E)** For the experiment described in (D), the net reduction in uncertainty of the EMT phenotype based on pERK, pc-Jun, or both signals is plotted. n = 4, one-way ANOVA with Tukey’s multiple comparisons test. **F)** PDX 395 cells that were treated with DMSO or 10 ng/mL 5-FU were treated with DMSO, 20 nM trametinib, 10 μM SP600125, or both inhibitors for 3 days. Representative 4i images of the indicated proteins, n = 3. **G)** For the experiment described in (F), the mosaic plot displays percentages of cells in each EMT phenotype bin. n = 3, Chi-squared test. **H)** HPAF-II cells were treated with 10 ng/mL gemcitabine or 10 ng/mL 5-FU for 48 hr followed by a washout with complete media for 72 hr. RPMI and DMSO were used as controls for gemcitabine or 5-FU, respectively. Representative live-cell fluorescence images of TO-PRO-3 (top) and paired fixed immunofluorescence images of vimentin (bottom), n = 3. **I)** For the experiment described in (H), the mosaic plot displays vimentin positivity in TO-PRO-3-negative cells. Indicated n = number of cells pooled across 3 biological replicates, Chi-squared test. **J)** HPAF-II cells that were treated with or without 1 ng/mL gemcitabine were treated with DMSO or the indicated concentrations of trametinib for 3 days. Representative immunofluorescence images of pERK and vimentin, n = 3. **K)** For the experiment described in (J), pERK intensity is plotted. n = 3, one-way ANOVA with Tukey’s multiple comparisons test. **L)** For the experiment described in (J), vimentin positivity is plotted. N = 3, one-way ANOVA with Tukey’s multiple comparisons test. For all panels *p <0.05, and error bars represent the standard error of the mean. Scale bars indicate 300 μm.

**Supplemental Figure S7.**
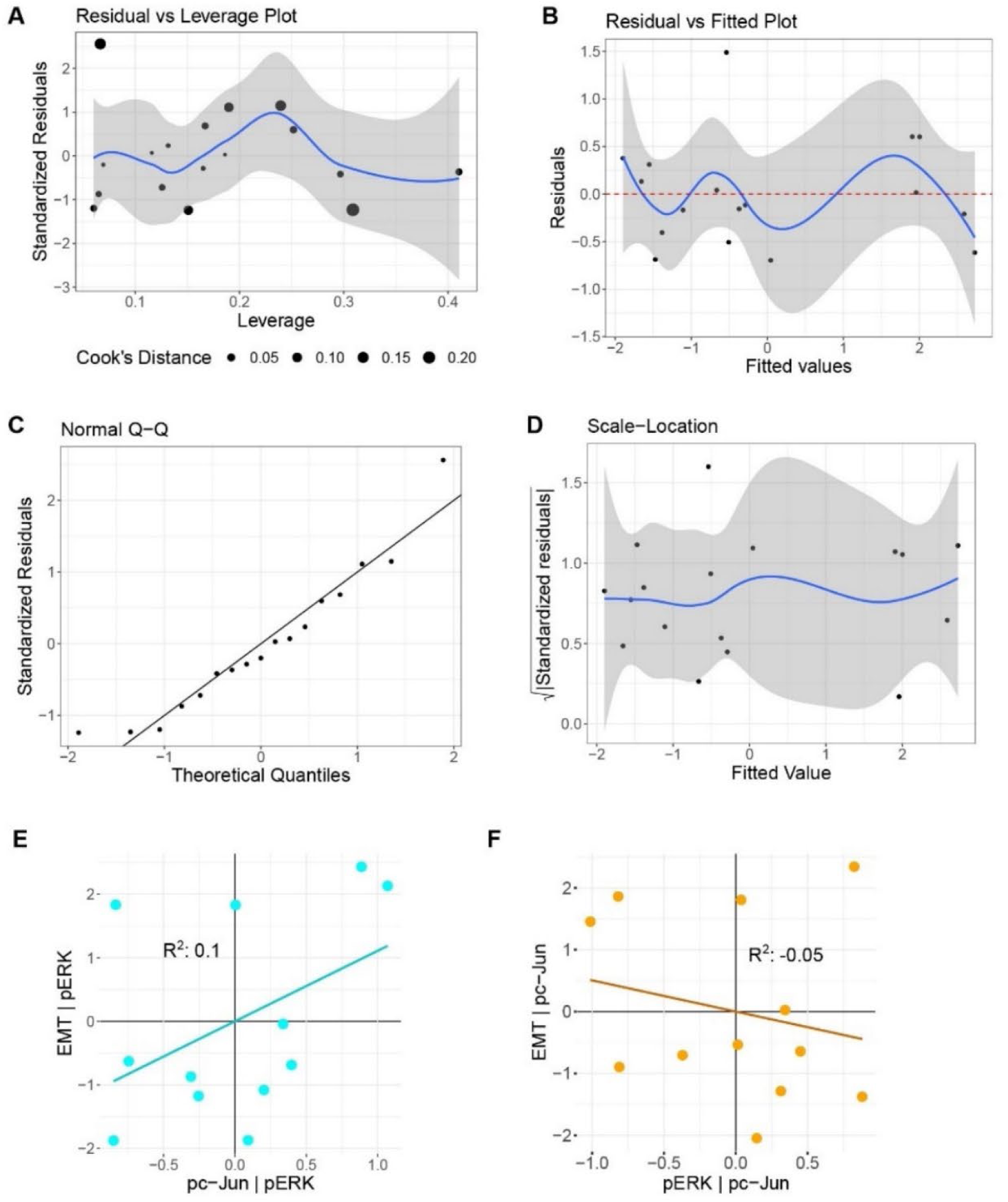
Population-level model meets regression model assumptions, and simulated bulk immunofluorescence data misses the role of ERK. For the population-level model of signaling and EMT based on western blot measurements described in Figure 5: **A)** A residuals-versus-leverage plot shows one outlier, which was removed from the model; **B)** A plot of fitted values versus residuals plot shows linearity; **C)** QQ plot shows normality; and **D)** scale-location plot shows homoscedasticity. For a model based on aggregated immunofluorescence data from Figure 1: **E)** Partial regression of EMT with pc-Jun, controlling for pERK, is shown. n = 3, linear least squares regression; **F)** Partial regression of EMT and pERK, controlling for pc-Jun, is shown. n = 3, linear least squares regression. For all panels *p < 0.05.

**Supplemental Figure S8.**
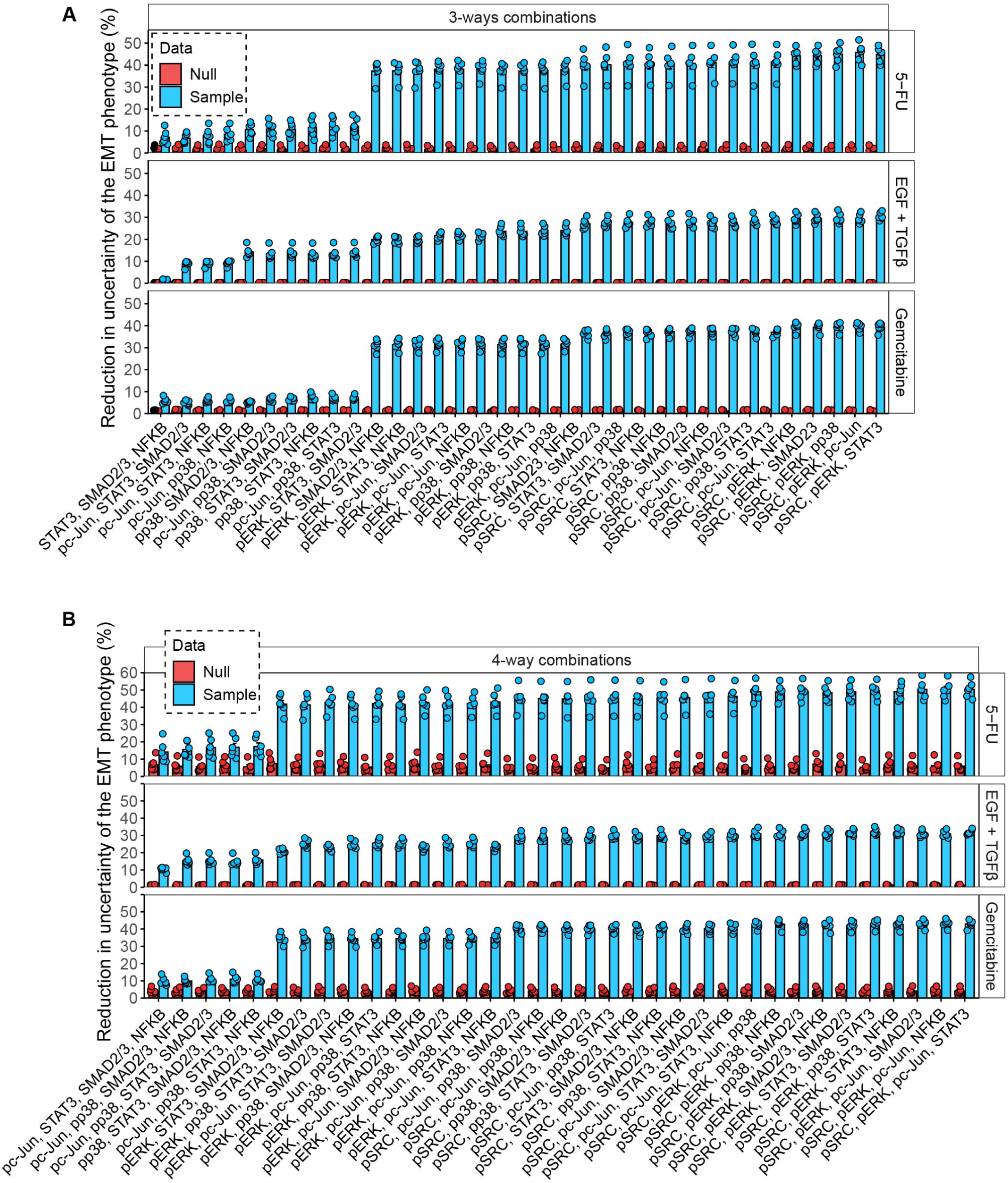
Extended mutual information analysis. HPAF-II cells were treated and subjected to 4i as described in Figure 6. The net percent reduction in uncertainty of the EMT phenotype from the indicated 3-way combinations of signals **(A)** or 4-way combinations of signals **(B)** are displayed. “Null” indicates the result for the shuffled null distribution. n = 6, error bars indicate standard error of the mean.

